# Bacterial Extracellular Vesicles from *Chromobacterium subtsugae* and *Bacillus thuringiensis* as Cell-Free Bioinsecticidal Nanocarriers Against the Soybean Pest *Euschistus heros*

**DOI:** 10.64898/2026.08.07.743497

**Authors:** Maria Eduarda Cimi, Daiane Gonzaga Ribeiro, Yasmim Omendes Nascimento, Micaela Cristiane Gomes dos Reis, Bruna Beatriz da Silva Ribeiro, Eliane Luiz de Freitas, Rodrigo Mauricio Marinsek Sales, Caroline Costa Lessa, Rosiane Andrade da Costa, Marcelo Tavares de Castro, Marina Arantes Radicchi, Sonia Nair Bao, Wagner Fontes, Rinaldo Wellerson Pereira, Rose Gomes Monnerat Solon de Pontes, Maria Sueli S. Felipe, Getulio Pereira de Oliveira

**Affiliations:** Graduate Program in Genomic Sciences and Biotechnology, Universidade Católica de Brasília, Brasília, DF, Brazil. CEP 71966-700; Laboratory of Protein Chemistry and Biochemistry, Department of Cell Biology, University of Brasília, Brasília, DF, Brazil. CEP 70910-900; Laboratory of anatomy, Faculty of Medicine, Department of Morphology, University of Brasília, Brasília, DF, Brazil. CEP 70910-900; Laboratory of Entomology, Embrapa Cerrados, Brazilian Agricultural Research Corporation, Brasília, DF, Brazil. CEP: 73310-970; Microscopy and Microanalysis Laboratory, Department of Cell Biology, University of Brasília, Brasília, DF, Brazil. CEP 70910-900; Arbor Garden, Brasília, DF, Brazil. CEP: 73020-407

**Author notes:** These authors contributed equally to this manuscript and share senior authorship (Maria Sueli S. Felipe, Getulio Pereira de Oliveira).

**Keywords:** bacterial extracellular vesicles, bioinsecticide, *Euschistus heros*, *Chromobacterium subtsugae*, *Bacillus thuringiensis*, violacein

## Abstract

Bacterial extracellular vesicles (bEVs) are membrane-enclosed nanoparticles that transport bioactive cargo and mediate interactions between bacteria and their environment. Although bEVs are increasingly recognized as natural delivery systems, their potential application in plant pest biocontrol remains poorly explored. Here, we provide proof-of-concept evidence that isolated bEVs from two entomopathogenic bacteria, *Chromobacterium subtsugae* and *Bacillus thuringiensis* var. *kurstaki*, exert insecticidal activity against the soybean pest *Euschistus heros*. Isolated bEVs were characterized by tunable resistive pulse sensing, nano-flow cytometry, transmission electron microscopy, SDS-PAGE, MALDI-TOF mass spectrometry, and label-free quantitative proteomics. *C. subtsugae* bEVs displayed a proteome clearly remodeled relative to the soluble protein fraction, with enrichment of outer-membrane, secretion-associated, proteolytic, and membrane-active proteins. MALDI-TOF analysis detected a violacein-associated ion selectively in the *C. subtsugae* bEV fraction, supporting vesicular association of this hydrophobic bioactive metabolite. In survival assays, *C. subtsugae* bEVs strongly reduced *E. heros* nymph survival (HR = 4.0, p < 0.0001), whereas the corresponding soluble protein fraction was inactive (HR = 1.2, p = 0.50). In contrast, *B. thuringiensis* bEVs and soluble protein fractions produced similar moderate activity (both HR = 2.1), consistent with their largely overlapping proteomic profiles. Cry1Ab was detected mainly in the *B. thuringiensis* soluble fraction rather than selectively enriched in bEVs. Together, these findings support a multi-component cargo model in which *C. subtsugae* bEVs combine vesicle-associated violacein with enriched protein cargo, establishing bacterial EVs as promising natural nanocarriers for next-generation, cell-free bioinsecticides against Cry-resistant hemipteran pests such as *E. heros*.

## Introduction

Soybean (*Glycine max* L.) is one of the most important crops worldwide, with Brazil, the United States, and Argentina among the leading producers (USDA-WASDE, 2025). Despite advances in crop management, soybean productivity remains severely affected by insect pests (Pozebon et al., 2020), particularly by the Neotropical brown stink bug *Euschistus heros* (Fabricius, 1798) (Hemiptera: Pentatomidae), one of the most damaging sap-sucking pests in South America (Saldanha et al., 2024). By feeding on pods and seeds during reproductive stages, *E. heros* impairs seed development, reduces grain quality, and facilitates opportunistic infections (Tessmer et al., 2022).

Current control of *E. heros* relies largely on repeated applications of broad-spectrum synthetic insecticides. This strategy has contributed to resistance in field populations, including target-site mutations associated with cross-resistance to multiple insecticide classes (Cuenca et al., 2025), and has increased production costs and environmental pressure (Silva et al., 2023). Sustainable pest-management strategies that reduce dependence on synthetic pesticides are therefore urgently needed.

Microbial biopesticides represent an important alternative in integrated pest management. Among them, *Chromobacterium subtsugae* and *Bacillus thuringiensis* are particularly relevant because they produce bioactive molecules with insecticidal potential. *C. subtsugae* is a Gram-negative soil bacterium that produces violacein, a hydrophobic indole-derived metabolite associated with antimicrobial and insecticidal activity (Martin et al., 2007). In contrast, *B. thuringiensis* var. *kurstaki* HD1-1450 is a Gram-positive entomopathogenic bacterium known to produce Cry1Ab toxins, which are primarily active against lepidopteran insects through receptor-mediated disruption of the midgut epithelium (Bravo et al., 2007). However, how insecticidal molecules from these bacteria are naturally packaged, protected, and delivered to target insects remains incompletely understood.

Bacterial extracellular vesicles (bEVs) provide a plausible mechanism for such delivery. These nanoscale, membrane-enclosed particles are produced by bacteria across phyla and can transport proteins, lipids, nucleic acids, and secondary metabolites (Toyofuku et al., 2023). In Gram-negative bacteria, bEVs are primarily released as outer membrane vesicles (OMVs), whereas Gram-positive bacteria produce cytoplasmic membrane vesicles through mechanisms involving cell-wall remodeling (Briaud and Carroll, 2020). Because bEVs can selectively package bioactive cargo, they are increasingly recognized as natural nanocarriers rather than simple cellular debris (Kim et al., 2015).

The potential relevance of bEVs in agriculture is only beginning to emerge. EVs from plant-associated and soil-associated bacteria have been implicated in interkingdom communication, plant growth promotion, and resistance to phytopathogens (Zannis-Peyrot et al., 2025). In entomopathogenic systems, bEVs may similarly transport enzymes, toxins, or metabolites that can disrupt disrupting insect physiology (Remans et al., 2025). Notably, the *B. thuringiensis* vegetative insecticidal protein Vip3Aa has been reported to associate with membrane vesicles (Zhang et al., 2022), suggesting that vesicle-mediated transport may occur naturally in insecticidal bacteria.

Despite this potential, the composition and functional role of bEVs from entomopathogenic bacteria remain poorly understood. An early study reported intrinsic insecticidal activity in purified OMVs from *Xenorhabdus nematophilus* (Khandelwal and Banerjee-Bhatnagar, 2003), but this observation has not been broadly extended to other bioinsecticidal bacteria. To date, no study has examined whether bEVs from *C. subtsugae* and *B. thuringiensis* contribute to insecticidal activity against *E. heros*, a hemipteran pest that is poorly targeted by conventional Bt Cry toxins due to limited toxin processing and receptor interactions in the gut (Chougule and Bonning, 2012).

Here, we hypothesized that extracellular vesicles produced by entomopathogenic bacteria package bioactive insecticidal cargo and contribute to insect mortality. To test this, we isolated bEVs and corresponding soluble protein fractions from *C. subtsugae* and *B. thuringiensis* var. *kurstaki*, characterized their biophysical and ultrastructural properties, profiled their proteomes by label-free quantitative mass spectrometry, performed targeted MALDI-TOF MS analysis of violacein, and evaluated their insecticidal activity against second-instar *E. heros* nymphs. This study provides evidence that bacterial EVs can function as natural carriers of insecticidal cargo and supports their further exploration as sustainable, non-replicative platforms for pest management.

## Material and methods

### Reagents

Dulbecco’s phosphate-buffered saline (dPBS; cat. no. 14190250) and CellMask™ Deep Red Plasma Membrane Stain (cat. no. C10046) were obtained from Thermo Fisher Scientific (Waltham, MA, USA).

### Bacterial strains, identification, and culture conditions

The *Chromobacterium subtsugae* and *Bacillus thuringiensis* var. *kurstaki* strains used in this study were maintained at −80 °C in 16% (v/v) glycerol.

Preliminary strain identity and purity were assessed by colony morphology on Tryptic Soy Agar (TSA). *B. thuringiensis* produced white, unpigmented colonies, whereas *C. subtsugae* displayed the characteristic purple pigmentation associated with violacein biosynthesis (**Supplementary Figure 1**).

For bEV production, seed cultures prepared from single colonies were used to inoculate stainless-steel bioreactors with working volumes of 5–50 L. Cultures were grown under controlled aeration, agitation, temperature, pH, and dissolved oxygen until stationary phase. *C. subtsugae* was cultured at 28 °C in a medium with a carbon-to-nitrogen (C/N) ratio of 15:1, and *B. thuringiensis* was cultured at 30 °C in a medium with a C/N ratio of 8:1. At the end of fermentation, cultures were transferred into sterile production bags, and bEVs were purified from the finalized formulations after biomass removal and sequential clarification. bEVs were purified from four independent fermentation batches per species.

### Bacterial extracellular vesicle isolation

Bacterial extracellular vesicles (bEVs) were isolated from culture supernatants by sequential clarification, ultrafiltration, and size-exclusion chromatography (SEC). Because *C. subtsugae* is Gram-negative and *B. thuringiensis* is Gram-positive, their vesicles are referred to as outer membrane vesicles (OMVs) and cytoplasmic membrane vesicles (CMVs), respectively; bEV is used throughout as an umbrella term (Welsh et al., 2024).

Culture supernatants were clarified by differential centrifugation at 300 × *g* for 10 min, 2,000 × *g* for 20 min, and 7,500 × *g* for 30 min to remove intact cells, large cellular debris, and residual large particles. Clarified supernatants were filtered through 0.22 µm syringe filters (Millipore, USA) and concentrated to 500 µL using Amicon® Ultra-15 centrifugal filter units with a 3 kDa molecular weight cutoff (MWCO; Merck Millipore, Billerica, MA, USA).

Concentrated samples were loaded onto qEVoriginal 35 nm SEC columns (Izon Science, Christchurch, New Zealand) pre-equilibrated with dPBS. The initial 2.5 mL void volume was collected as a control; the subsequent 2 mL bEV-enriched fraction was collected separately; the following 1.5 mL was discarded to minimize carryover; and the next 6 mL of later-eluting soluble protein-enriched fractions were collected for comparative analysis. The bEV- enriched fraction was further concentrated to 100 µL using Amicon® Ultra-2 centrifugal filter units with a 100 kDa MWCO. Void, bEV-enriched, and soluble protein fractions were stored at 4 °C until downstream characterization.

### Identification of violacein by MALDI-TOF mass spectrometry

Secondary metabolites in bEV-enriched and soluble protein fractions from *C. subtsugae* and *B. thuringiensis* were analyzed by MALDI-TOF mass spectrometry on an Autoflex Speed instrument (Bruker Daltonics, Bremen, Germany) equipped with a SmartBeam laser and controlled by FlexControl 3.0. Spectra were acquired in MS (TOF) mode only. External calibration was performed before each acquisition session using Peptide Calibration Standard II (Bruker Daltonics). Because this standard does not extend into the low-mass region, calibration in the range relevant to violacein detection was performed using the well-characterized ions of the CHCA matrix itself as calibrants.

Samples were mixed with α-cyano-4-hydroxycinnamic acid matrix (CHCA; 10 mg/mL in 50% acetonitrile, 49.7% ultrapure water, and 0.3% trifluoroacetic acid) at a 1:3 sample:matrix ratio, spotted onto a Bruker MTP Small Anchor 384 target plate, and air-dried for 24 h. Spectra were acquired in positive reflector mode using 500 laser shots per spot and triplicate technical replicates. Baseline subtraction, Savitzky–Golay smoothing, and SNAP peak detection with a signal-to-noise threshold >3 were applied uniformly in FlexAnalysis v3.4. Mass fingerprints were compared by m/z distribution and peak intensity, with particular attention to indole-derived metabolites associated with *C. subtsugae*, including violacein, expected at approximately m/z 344 as [M+H]⁺, and deoxyviolacein, expected at approximately m/z 328.1. Analyses were qualitative, and no quantitative inference was made from MALDI-TOF peak intensities.

### bEV characterization by tunable resistive pulse sensing

Purified bEVs were analyzed by tunable resistive pulse sensing (TRPS) using a qNano system (Izon Science) to determine particle size distribution and concentration. Samples were gently vortexed and diluted in sterile 1× PBS to approximately 1 × 10⁸ particles/mL. Measurements were performed using NP150 nanopores with an expected measurement range of approximately 70–300 nm. Acquisition settings were: stretch, 49.99 mm; voltage, 0.78 V; applied pressure, 15 cm H₂O; and baseline current, 112–121 nA. Calibration was performed immediately before each acquisition session using CPC200 calibration beads (203 ± 5 nm; Izon Science) under identical settings. At least 500 valid blockade events were recorded per sample. Data were processed in Izon Control Suite v3.3 to calculate particle size distribution and concentration.

### bEV characterization by nano-flow cytometry

Purified bEVs were analyzed on a Beckman Coulter CytoFLEX S equipped with violet side-scatter detection from the 405 nm laser. Sterile 0.22 µm-filtered PBS was used as sheath fluid and negative control. PBS baseline was maintained at 1,200–1,500 events/s, and bEV samples were diluted to acquisition rates of approximately 3,000–8,000 events/s to minimize coincident detection.

Instrument calibration was performed using NanoVis low and high polystyrene reference beads (Beckman Coulter; 80–600 nm). Calibration curves were analyzed in FCMPASS to convert side-scatter intensity into estimated particle diameter using a biological nanoparticle refractive index range of 1.37–1.40.

For fluorescence detection, bEVs were stained with CellMask™ Deep Red Plasma Membrane Stain for 30 min at room temperature in the dark and washed by SEC followed by ultrafiltration to remove unbound dye and micellar aggregates. Dye-only controls lacking bEVs were processed in parallel. Samples were acquired in Nano Mode at low flow rate (∼10 µL/min), with Violet SSC trigger threshold set at 1,775. CellMask Deep Red-stained vesicles were excited at 638 nm, and fluorescence emission was detected at 666 nm. Each sample was acquired for 2 min. Particle concentration was calculated by subtracting PBS background events, dividing by the gated acquisition volume, and multiplying by the dilution factor. Data were analyzed in FlowJo v10.8.1 using a 2-min time gate and debris-exclusion gates defined from PBS background and bead calibration.

### bEV characterization by fluorescence microscopy

Purified *C. subtsugae* bEVs were stained with CellMask™ Deep Red Plasma Membrane Stain for 30 min at room temperature in the dark and washed by SEC followed by ultrafiltration. Uninoculated Bug Medium® processed through the complete bEV isolation workflow and stained in parallel was used as a negative control.

For imaging, 5 µL of each stained preparation was mounted on glass slides under coverslips and examined using a Zeiss Axio Imager.M2 microscope with a 100× objective and Texas Red filter set. Although this filter set provides suboptimal excitation for CellMask Deep Red, sufficient signal was obtained to confirm membrane labeling of the purified bEV fraction. Images were displayed in pseudocolor.

### bEV characterization by transmission electron microscopy

Purified bEV preparations, void fractions, and soluble protein fractions were analyzed by negative-stain transmission electron microscopy (TEM). For each sample, 5 µL was adsorbed onto carbon-coated 200-mesh copper grids (Electron Microscopy Sciences, Hatfield, PA, USA) for 15 min at room temperature. Grids were washed with ultrapure water, negatively stained with 3% uranyl acetate for 1 min, and air-dried for 24 h.

Images were acquired on a JEOL JEM-1010 microscope (Tokyo, Japan) operated at 80 kV and equipped with a Gatan Orius SC1000 CCD camera. At least five micrographs were acquired per sample. Representative images were processed in ImageJ (NIH, USA), and scale bars were applied consistently.

### Silver-stained SDS-PAGE of bEVs

Protein concentrations in purified bEV and soluble protein fractions were determined using the Pierce™ BCA Protein Assay Kit (QuantPro; Thermo Fisher Scientific). For each lane, 5 µg of total protein was mixed with 2× Laemmli buffer containing β-mercaptoethanol at a final concentration of 5%, heated at 99 °C for 10 min, and loaded onto in-house 15% Tris-Glycine polyacrylamide gels.

Electrophoresis was performed using the Bio-Rad Mini-PROTEAN system at 150 V for approximately 45 min. Gels were silver-stained manually using standard fixation, sensitization, silver nitrate incubation, development, and stop steps, and were photographed immediately after staining.

### Label-free quantitative proteomics Protein digestion and peptide desalting

Protein digestion was performed by filter-aided sample preparation (FASP) as described by (Wiśniewski et al., 2009), with minor adaptations. Three independent biological replicates of purified bEV-enriched fractions and soluble protein fractions were prepared from each bacterial species, yielding 12 samples in total.

For each sample, 100 µg of total protein was solubilized in 8 M urea, pH 8.5, and loaded onto Vivacon 500 centrifugal filter units with a 30 kDa MWCO (Sartorius Stedim, CA, USA). Proteins were reduced with 0.1 M DTT, alkylated with 20 mM iodoacetamide, washed with 20 mM triethylammonium bicarbonate, and digested with sequencing-grade trypsin at an enzyme:protein ratio of 1:100 for 16 h at 37 °C. Peptides were acidified with 10% trifluoroacetic acid, dried, resuspended in 0.1% formic acid, desalted using in-house C18/Poros Oligo R2 microcolumns as described by (Tahir et al., 2020), quantified by Qubit, and resuspended in 0.1% formic acid before LC-MS/MS analysis.

### Liquid chromatography and mass spectrometry

Peptides were separated on a Dionex Ultimate 3000 RSLCnano system (Thermo Scientific), loaded onto a trap column, and resolved on a 24 cm × 75 µm analytical column using a 155 min linear gradient from 2% to 35% solvent B. Solvent A consisted of 0.1% formic acid and 2% acetonitrile, and solvent B consisted of 0.1% formic acid and 80% acetonitrile. Peptides were analyzed on an Orbitrap Elite mass spectrometer (Thermo Scientific) operated in data-dependent acquisition mode. MS1 spectra were acquired at 120,000 resolution across the *m/z* 300–1650 range, and the top 15 ions above an intensity threshold of 3,000 were selected for MS/MS fragmentation.

### Protein identification and quantification

Label-free feature detection, alignment, and quantification were performed in Progenesis QI for Proteomics v1.0 (Nonlinear Dynamics), with a minimum alignment score of 70% and features restricted to charge states 2–7.

Protein identification was performed in PEAKS Studio v7.0 (Bioinformatics Solutions Inc.) using de novo sequencing combined with database searching against restricted UniProt reference proteomes for *C. subtsugae* (taxon ID 251747) and *B. thuringiensis* (taxon ID 1428), using the March 2026 database release. Search parameters included a precursor tolerance of 10 ppm, fragment tolerance of 0.05 Da, up to two missed cleavages, carbamidomethylation of cysteine as a fixed modification, and methionine oxidation as a variable modification. SPIDER was used to identify unexpected modifications. Protein identifications were accepted at a protein-level false discovery rate below 1%.

### Differential abundance and bioinformatics analysis

Quantified protein abundances were analyzed in R using the DEP2 package (Feng et al., 2023). The workflow included variance-stabilizing normalization, minimal-probability imputation of missing values, and differential-abundance testing using moderated *t*-tests implemented in limma, with Benjamini–Hochberg correction.

Proteins with an adjusted *p*-value < 0.05 and |log₂(fold change)| > 1 were considered differentially abundant. This threshold was prespecified and applied identically to both species. The contrast was defined as soluble protein fraction versus bEV fraction; therefore, negative log₂(fold change) values indicate enrichment in the bEV fraction, whereas positive values indicate enrichment in the soluble protein fraction.

For *B. thuringiensis*, an exploratory relaxed threshold of adjusted *p*-value < 0.10 was additionally applied to nominate candidate vesicle-associated proteins for Gene Ontology over-representation analysis. Proteins identified only under this threshold were treated as exploratory candidates rather than statistically significant differences. Hierarchical clustering, principal component analysis, Venn analysis of fraction-exclusive proteins, and Gene Ontology over-representation analyses were performed within the same analytical framework, using UniProt functional annotations.

### Evaluation of bacterial bEVs and soluble protein fractions as bioinsecticides against ***Euschistus heros***

#### Insect rearing and maintenance

*Euschistus heros* eggs were obtained from a commercial rearing facility. Egg masses were collected every 48 h and placed in sterile 9 cm Petri dishes lined with moistened filter paper and sealed with Parafilm®. First-instar nymphs were maintained for 48 h with humidity provided by a moistened cotton pad and without supplementary feeding.

From the second instar onward, nymphs were transferred to cylindrical rearing containers in groups of 15–20 individuals at a similar developmental stage. Containers were maintained at 25.8 ± 3.2 °C, 55.2 ± 13.8% relative humidity, and a 12 h light:12 h dark photoperiod. Fresh green bean pods (*Phaseolus vulgaris* L.) were provided ad libitum and replaced every 48 h.

#### Bioassay design and procedures

Bioassays were performed using newly molted second-instar nymphs (<12 h post-ecdysis) to ensure synchronized developmental stage and comparable physiological status. The assay was adapted from the artificial feeding system described by (Castro et al., 2025). Each treatment initially comprised 28 nymphs distributed across four replicate 50 mL Falcon® tubes, with seven nymphs per tube. Tubes contained a cotton support at the base and were sealed with Parafilm® to form a feeding sachet. Nymphs were acclimated for 48 h with 300 µL of sterile bean-based artificial diet. Only individuals alive and actively feeding at treatment onset were retained for survival analysis, resulting in final sample sizes of 19–26 nymphs per treatment.

At treatment onset (*t* = 0), 100 µL of each test preparation was added to the existing 300 µL of bean-based diet, yielding a final diet:treatment ratio of 3:1 (v/v). Treatments were: sterile diet alone as negative control; Grandevo® at 350 µg of formulated product per dose as positive control; *C. subtsugae* whole culture supernatant, 100 µL per dose; purified *C. subtsugae* bEVs, 6 × 10¹¹ vesicles and approximately 284 µg protein per dose; purified *B. thuringiensis* bEVs, 6 × 10¹¹ vesicles and approximately 136 µg protein per dose; and soluble protein fractions from *C. subtsugae* or *B. thuringiensis*, 150 µg total protein per dose. Nymphs were maintained for 168 h under bioassay conditions of 22 °C, 45% relative humidity, and a 12 h light:12 h dark photoperiod. Mortality was recorded every 24 h. Dead nymphs were defined as individuals that showed no response to gentle tactile stimulation and remained immobile for more than 1 min. Survival data reported in the main analysis derive from a single representative bioassay; additional independent bioassays performed under identical conditions yielded consistent results.

### Statistical analysis

Statistical analyses were performed in GraphPad Prism v8.1 (GraphPad Software, San Diego, CA, USA). For TRPS and nano-flow cytometry, pooled particle size distributions were compared between species using chi-square tests of homogeneity after binning the data into 10 nm intervals. Mean and modal particle diameters, as well as particle concentrations, were compared between species using unpaired two-tailed *t*-tests with Welch’s correction. Survival data from the 168 h bioassay were analyzed using the log-rank Mantel–Cox test for pairwise comparisons with the negative control. Multiple comparisons were adjusted using the Holm–Šidák correction. For each treatment, hazard ratios relative to the negative control were estimated using the Mantel–Haenszel method and reported with 95% confidence intervals and corresponding adjusted log-rank *p*-values. Results are expressed as mean ± SD with 95% confidence intervals where applicable. Statistical significance was set at α = 0.05 and reported as ns, *p* ≥ 0.05; *p* < 0.05; *p* < 0.01; *p* < 0.001; and *p* < 0.0001.

## Results

### Isolation and characterization of bEVs

Bacterial cultures were processed by sequential clarification, ultrafiltration, and size-exclusion chromatography (SEC) to obtain two operationally distinct preparations from each species: bEV-enriched fractions and soluble protein-enriched fractions. These preparations were then analyzed using complementary biophysical, biochemical, molecular, and functional approaches, including TRPS, nano-flow cytometry, TEM, SDS-PAGE, MALDI-TOF mass spectrometry, label-free quantitative proteomics, and insect survival bioassays. An overview of the experimental workflow is presented in **Figure 1**.

**Figure 1.**
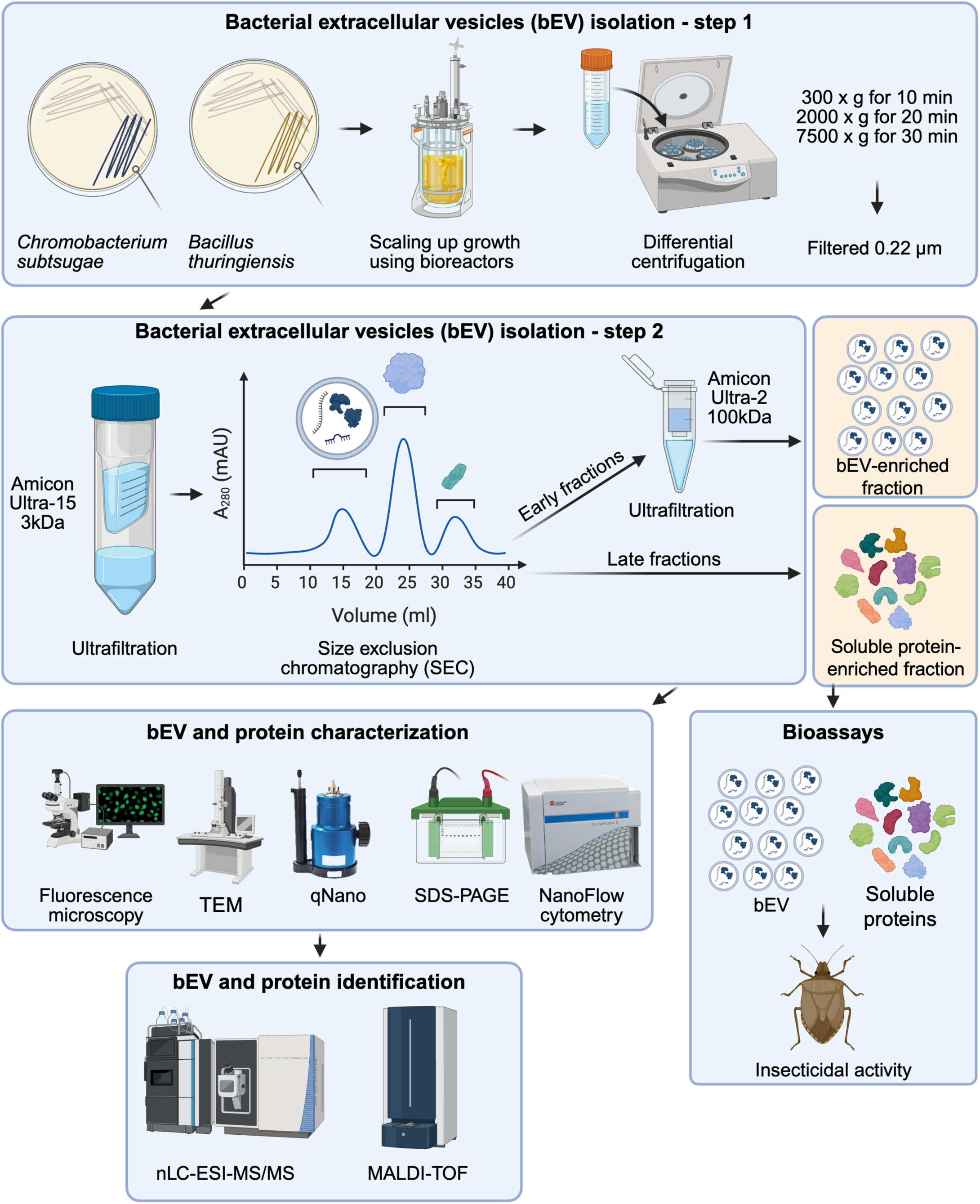
Experimental workflow for isolation, characterization, molecular profiling, and bioassay evaluation of bacterial extracellular vesicles (bEVs). Bacterial strains (*Chromobacterium subtsugae* and *Bacillus thuringiensis*) were cultured and expanded in bioreactors. Step 1: clarified supernatants were processed through sequential differential centrifugation (300 × g for 10 min to remove intact cells, 2,000 × g for 20 min to eliminate large cellular debris, and 7,500 × g for 30 min to remove residual particles), followed by 0.22 μm filtration to remove remaining bacterial cells, spores, and large contaminants. Step 2: filtered supernatants were concentrated using 3 kDa ultrafiltration units and fractionated by size-exclusion chromatography (SEC). Early-eluting fractions enriched in bEVs were collected separately from later-eluting fractions enriched in soluble proteins, and the bEV-enriched fraction was further concentrated by 100 kDa ultrafiltration, yielding two purified materials: bEV-enriched fractions and soluble protein-enriched fractions. Characterization: both fractions underwent multimodal biophysical and biochemical analyses, including fluorescence microscopy (CellMask labeling), transmission electron microscopy (TEM; negative staining), tunable resistive pulse sensing (qNano; particle size and concentration), nano-flow cytometry (CytoFLEX S; size distribution and membrane-lipid content), and silver-stained SDS-PAGE (protein profile). Molecular profiling was conducted by matrix-assisted laser desorption/ionization–time-of-flight mass spectrometry (MALDI-TOF MS) for metabolite identification and by nLC-ESI-MS/MS for quantitative proteomics. Bioassays: purified bEVs and soluble protein fractions were incorporated into sterile artificial insect diet at defined doses and administered to *Euschistus heros* second-instar nymphs. Insecticidal activity was assessed by 168-hour survival assays against negative (sterile diet) and positive (Grandevo®) controls.

SEC separated early-eluting bEV-enriched fractions from later-eluting soluble protein fractions for both bacterial species. After SEC and concentration by 100 kDa ultrafiltration, the *C. subtsugae* bEV-enriched fraction retained a characteristic purple coloration, consistent with the presence of violacein-associated material in the vesicle fraction (**Figure 2A**). In contrast, *B. thuringiensis* bEV preparations remained colorless before and after ultrafiltration, as expected for a species that does not naturally produce violacein-like pigments (**Figure 2B**). Soluble protein fractions from both species displayed yellowish-brown coloration typical of protein-rich bacterial culture material, while void and flow-through fractions were collected as separation controls. These visual differences provided initial evidence that pigment-associated metabolites preferentially partitioned into the *C. subtsugae* bEV fraction.

**Figure 2.**
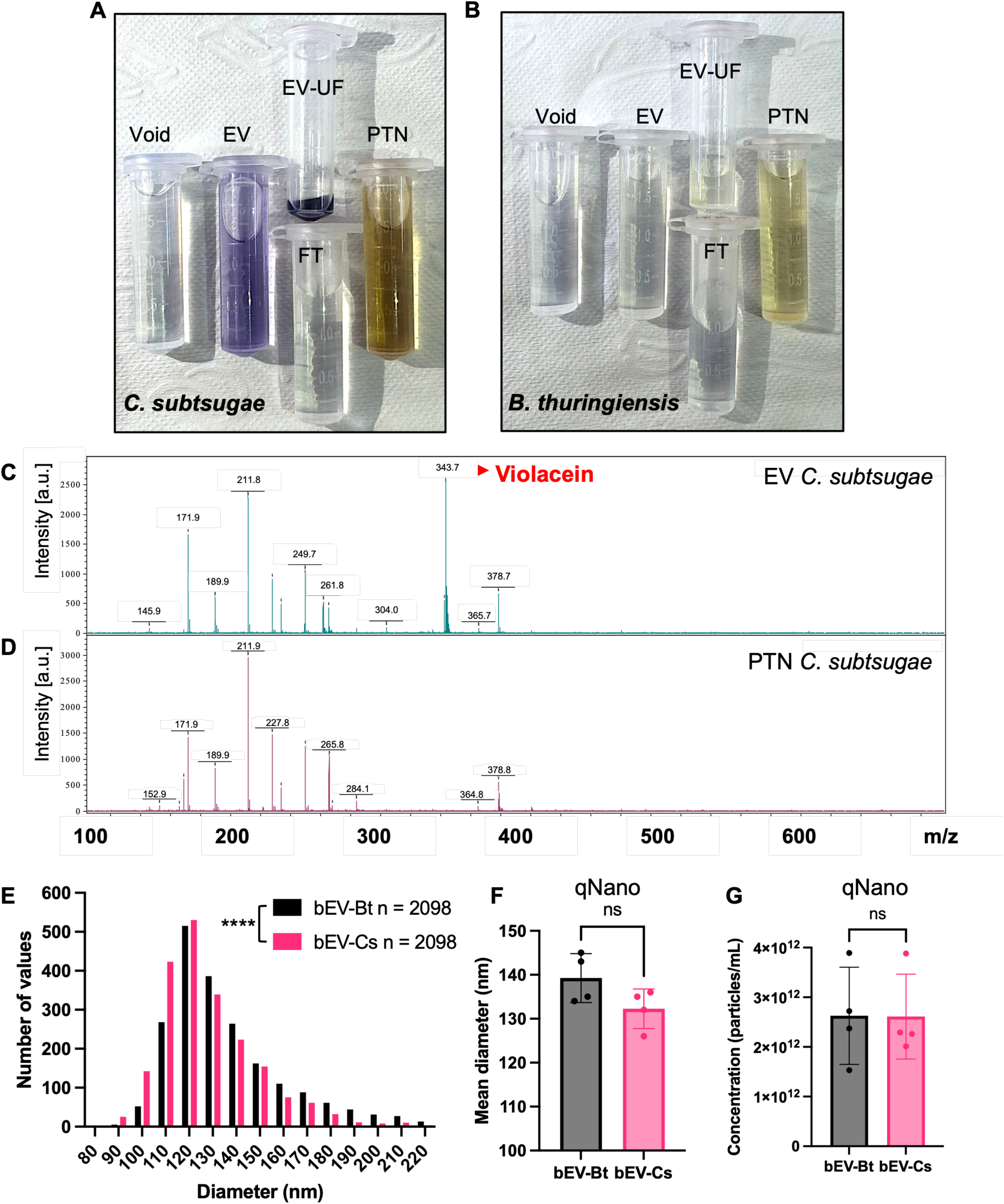
Purification and characterization of extracellular vesicles (bEVs) from Chromobacterium subtsugae *and* Bacillus thuringiensis. (A) Size-exclusion chromatography (SEC) and ultrafiltration fractions from *C. subtsugae*: void volume (Void), EV-enriched fraction (EV), flow-through (FT), concentrated EV-enriched retentate (EV-UF), and soluble protein fraction (PTN). (B) SEC and ultrafiltration fractions from *B. thuringiensis*: void volume (Void), EV-enriched fraction (EV), flow-through (FT), concentrated EV-enriched retentate (EV-UF), and soluble protein fraction (PTN). (C) MALDI-TOF mass spectrum of the *C. subtsugae* EV-enriched fraction, with annotated m/z values for major peaks; the peak at m/z 344 corresponds to violacein. (D) MALDI-TOF mass spectrum of the *C. subtsugae* soluble protein (PTN) fraction, with annotated m/z values for major peaks. (E) Vesicle size-distribution profiles by tunable resistive pulse sensing (TRPS) for *B. thuringiensis* bEVs (bEV-Bt, n = 2,098 particles) and *C. subtsugae* bEVs (bEV-Cs, n = 2,098 particles). (F) Mean vesicle diameter determined by TRPS for bEV-Bt and bEV-Cs (mean ± SD; n = 4 preparations; ns, not significant). (G) Particle concentration determined by TRPS for bEV-Bt and bEV-Cs (mean ± SD; n = 4 preparations; ns, not significant). [Figure image: relabel the violacein peak from m/z 343.7 to m/z 344 (or “[M+H]⁺ 344”) to match the text.]

### MALDI-TOF mass spectrometry identifies violacein-associated ions in *C. subtsugae* bEVs

To determine whether violacein was associated with the *C. subtsugae* bEV fraction, purified bEVs and corresponding soluble protein fractions were analyzed by MALDI-TOF mass spectrometry. A certified violacein analytical standard produced an ion at m/z 343.7 under the acquisition conditions used, close to the theoretical protonated violacein ion [M+H]⁺ at m/z 344.1. Because the same acquisition and calibration conditions were applied to standards and samples, identification was based on direct correspondence with the analytical standard rather than on absolute mass accuracy.

Appropriate controls, including MALDI matrix blanks and uninoculated culture media processed for both bacterial species, were used to evaluate background ions and specificity (**Supplementary Figure 2A,B**). *B. thuringiensis* bEV and soluble protein fractions did not show an ion matching the violacein standard, and their spectra were dominated by matrix-associated background peaks (**Supplementary Figure 2C–F**).

In contrast, the *C. subtsugae* bEV fraction showed a prominent ion at m/z 343.7, matching the violacein standard under the same analytical conditions (**Figure 2C**). This ion was not detected in the corresponding *C. subtsugae* soluble protein fraction, which showed primarily matrix-associated peaks and additional low-intensity ions not attributable to violacein (**Figure 2D**). Additional ions at m/z 249.7, 261.8, and 304.0 were observed in the bEV fraction and may represent violacein-related fragments or associated metabolites; however, these assignments were not confirmed by MS/MS and should therefore be interpreted cautiously. Together, these MALDI-TOF data show that a violacein-associated ion was detected in the *C. subtsugae* bEV-enriched fraction, but not in the matched soluble protein fraction or in *B. thuringiensis* preparations.

### TRPS reveals distinct vesicle size-distribution profiles between species

Tunable resistive pulse sensing was used to compare bEV size distributions and particle concentrations between *C. subtsugae* and *B. thuringiensis*. The two species showed significantly different size-distribution profiles (**Figure 2E**). *C. subtsugae* bEVs were more broadly distributed, with enrichment in smaller diameter bins and detectable particles extending into larger size ranges. In contrast, *B. thuringiensis* bEVs showed a more homogeneous distribution centered around an intermediate size range. This difference in distribution shape was significant by chi-square test of homogeneity (p < 0.0001; n = 2,098 particles per species).

Despite these distributional differences, mean vesicle diameter did not differ significantly between species, with values of approximately 132 nm for *C. subtsugae* bEVs and 139 nm for *B. thuringiensis* bEVs (**Figure 2F**). Final particle concentrations were also comparable between preparations, at approximately 2.9 × 10¹² and 2.8 × 10¹² particles/mL for *C. subtsugae* and *B. thuringiensis*, respectively (**Figure 2G**). Because fermentations were not normalized by equivalent cell densities and the two species were grown in different media, these values represent concentrations of the final purified preparations rather than relative vesicle productivity per bacterial cell.

### Nano-flow cytometry confirms vesicle detection and membrane labeling

Nano-flow cytometry provided an orthogonal assessment of particle detection, scatter- derived size estimation, concentration, and membrane labeling. Instrument calibration using NanoVis polystyrene beads and FCMPASS showed strong model fitting, with (R² = 0.9967), and supported scatter-based estimation of vesicle size within the detectable range (**Supplementary Figure 3A–D**). PBS controls established a lower detection limit of approximately 110 nm, and both bEV preparations showed unimodal scatter profiles above background (**Figure 3A**).

**Figure 3.**
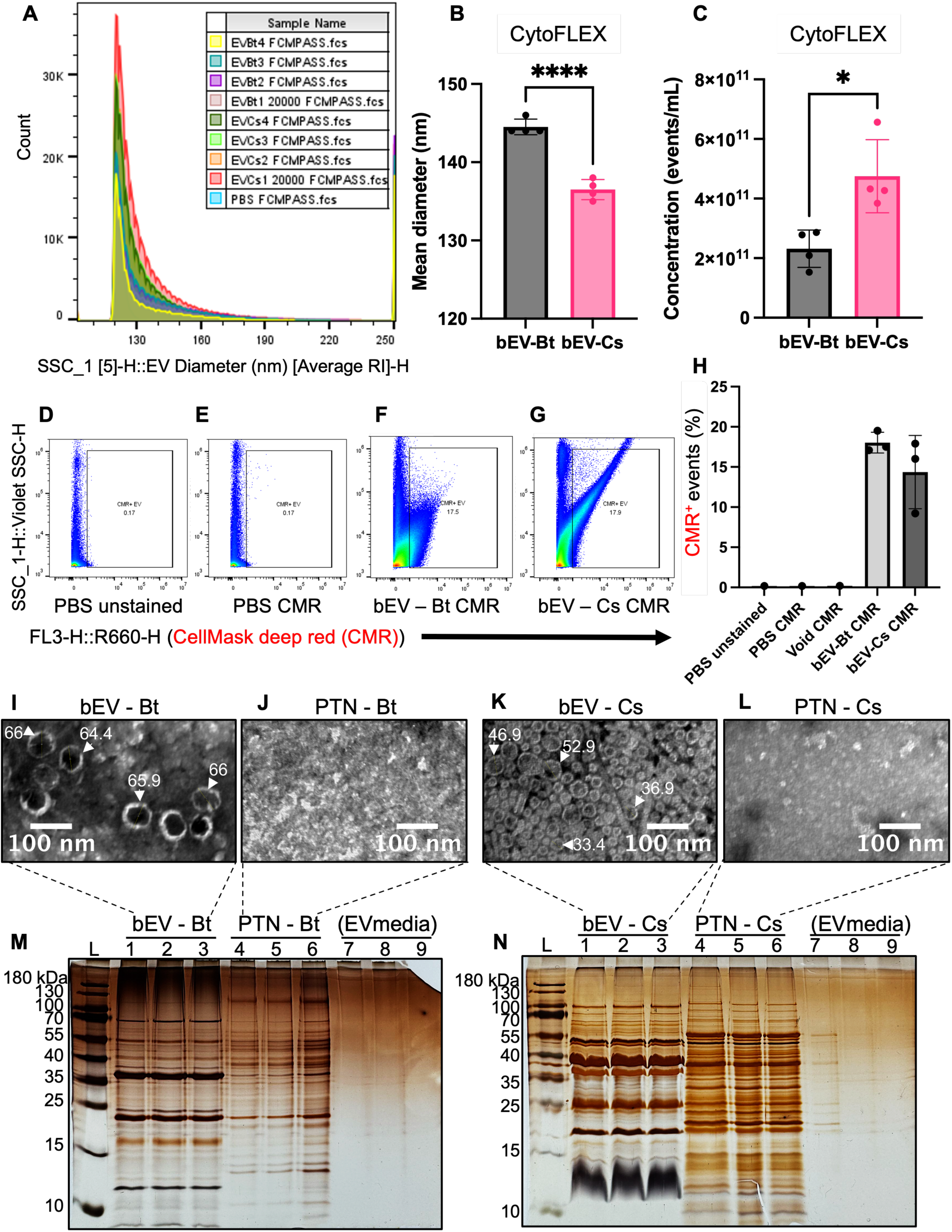
Biophysical and ultrastructural characterization of extracellular vesicles (bEVs) from *Chromobacterium subtsugae* and *Bacillus thuringiensis*. (A) Overlaid nano-flow cytometry side-scatter (SSC) histograms of purified bEV preparations from both species and PBS reference measurements. (B) Mean vesicle diameter for bEV-Bt and bEV-Cs by scatter-derived size estimation (nano- flow cytometry; mean ± SD; n = 4; ****p < 0.0001). (C) Particle concentration (events/mL) for bEV-Bt and bEV-Cs under identical acquisition parameters (mean ± SD; n = 4; *p < 0.05). (D) Nano-flow cytometry dot plot of PBS, unstained (gating control for CMR⁺ events). (E) Nano-flow cytometry dot plot of PBS incubated with CellMask Deep Red (CMR; dye-only control). (F) Nano-flow cytometry dot plot of *B. thuringiensis* bEVs stained with CellMask Deep Red. (G) Nano-flow cytometry dot plot of *C. subtsugae* bEVs stained with CellMask Deep Red. (H) Percentage of CMR⁺ events across conditions (PBS unstained, PBS + CMR, void + CMR, bEV-Bt + CMR, bEV-Cs + CMR); bars show mean ± SD. (I) Transmission electron microscopy (TEM) micrograph of the *B. thuringiensis* bEV-enriched fraction, with arrows indicating individual vesicles and their measured diameters (64.4–66 nm). Scale bar = 100 nm. (J) TEM micrograph of the *B. thuringiensis* soluble protein-enriched fraction. Scale bar = 100 nm. (K) TEM micrograph of the *C. subtsugae* bEV-enriched fraction, with arrows indicating individual vesicles and their measured diameters (33.4–52.9 nm). Scale bar = 100 nm. (L) TEM micrograph of the *C. subtsugae* soluble protein-enriched fraction. Scale bar = 100 nm. (M) Silver-stained SDS-PAGE of *B. thuringiensis* bEV-enriched fractions (lanes 1–3), soluble protein-enriched fractions (lanes 4–6), and EV-blank medium control (lanes 7–9). L, molecular weight ladder. (N) Silver-stained SDS-PAGE of *C. subtsugae* bEV-enriched fractions (lanes 1–3), soluble protein-enriched fractions (lanes 4–6), and EV-blank medium control (lanes 7–9). L, molecular weight ladder.

Scatter-derived size estimation indicated that *B. thuringiensis* bEVs were modestly but significantly larger than *C. subtsugae* bEVs, with mean diameters of approximately 145 nm and 137 nm, respectively (**Figure 3B**). Nano-flow cytometry also detected fewer *B. thuringiensis* bEV events than *C. subtsugae* bEV events under matched acquisition conditions (**Figure 3C**), suggesting differences in particle concentration, scattering behavior, or detection efficiency between species and platforms.

To assess membrane-associated labeling, bEVs were stained with CellMask Deep Red and washed by SEC followed by ultrafiltration to remove unbound dye and micellar aggregates. Fluorescence microscopy of stained *C. subtsugae* bEVs showed a dense field of fluorescent puncta, whereas uninoculated medium processed through the same workflow showed minimal signal (**Supplementary Figure 4**). Control experiments further showed that unwashed CellMask Deep Red produced substantial background fluorescence even at low dye concentrations, emphasizing the importance of post-staining SEC/UF washing (**Supplementary Figure 5**).

After washing, unstained PBS and PBS incubated with CellMask Deep Red showed minimal background signal, with 0.17% positivity in both controls (**Figure 3D,E**). In contrast, stained bEV samples showed clear membrane-associated fluorescence, with *B. thuringiensis* bEVs displaying approximately 18% CellMask-positive events and *C. subtsugae* bEVs approximately 14.3% CellMask-positive events (**Figure 3F,G**). Quantification across all conditions, including the stained void control, which showed low fluorescence positivity (0.20%), confirmed that positive signal was restricted to the bEV-containing samples (**Figure 3H**). These data support the presence of membrane-enclosed particles in the purified bEV fractions and confirm that the staining signal was not explained by free dye alone.

### TEM confirms vesicular ultrastructure and fraction enrichment

Negative-stain transmission electron microscopy directly visualized vesicle morphology. *B. thuringiensis* bEVs showed well-defined spherical or cup-shaped structures with clear membrane boundaries and relatively uniform morphology, with representative vesicles measuring approximately 64–66 nm in diameter (**Figure 3I**). The corresponding soluble protein fraction displayed mainly amorphous material with few organized vesicular structures (**Figure 3J**).

*C. subtsugae* bEVs showed higher particle density and greater morphological heterogeneity, including cup-shaped vesicular structures, apparent clusters, and representative diameters ranging from approximately 33 to 53 nm (**Figure 3K**). The corresponding soluble protein fraction contained sparse material and few vesicle-like structures (**Figure 3L**). As expected, TEM-derived diameters were smaller than those obtained by TRPS and nano-flow cytometry, consistent with the effects of dehydration and negative staining. Notably, *C. subtsugae* bEVs stored at 4 °C for 120 days retained recognizable vesicular morphology, suggesting structural stability under these storage conditions (**Supplementary Figure 6**).

### SDS-PAGE shows distinct protein profiles between bEV and soluble fractions

Silver-stained SDS-PAGE revealed distinct protein profiles between bEV-enriched and soluble protein fractions from both bacterial species. *B. thuringiensis* bEVs showed prominent bands mainly in the 30–55 kDa range, whereas the corresponding soluble protein fraction displayed a broader distribution of bands from approximately 15 to 100 kDa (**Figure 3M**). The EV-blank medium control showed minimal protein signal.

*C. subtsugae* bEVs displayed a more complex and protein-rich banding pattern, with intense bands across a broad molecular weight range from approximately 10 to 180 kDa (**Figure 3N**). A prominent pigmented band was observed near 12 kDa in the *C. subtsugae* bEV fraction, whereas this feature was absent from the corresponding soluble protein fraction. Given the much lower molecular mass of violacein itself, this band should not be interpreted as free violacein; rather, it may represent pigment-associated material co-migrating with vesicle-associated proteins, lipids, or membrane fragments. Overall, SDS-PAGE supported biochemical differences between bEV-enriched and soluble fractions and further indicated species-specific cargo composition.

### *B. thuringiensis* bEV and soluble protein fractions show highly similar proteomic profiles

Label-free quantitative proteomics was performed on *B. thuringiensis* bEV-enriched and soluble protein fractions using three independent biological replicates per fraction. Across replicates, bEV preparations yielded 125, 141, and 142 protein identifications, whereas soluble protein fractions yielded 152, 152, and 151 identifications (**Figure 4A**).

**Figure 4.**
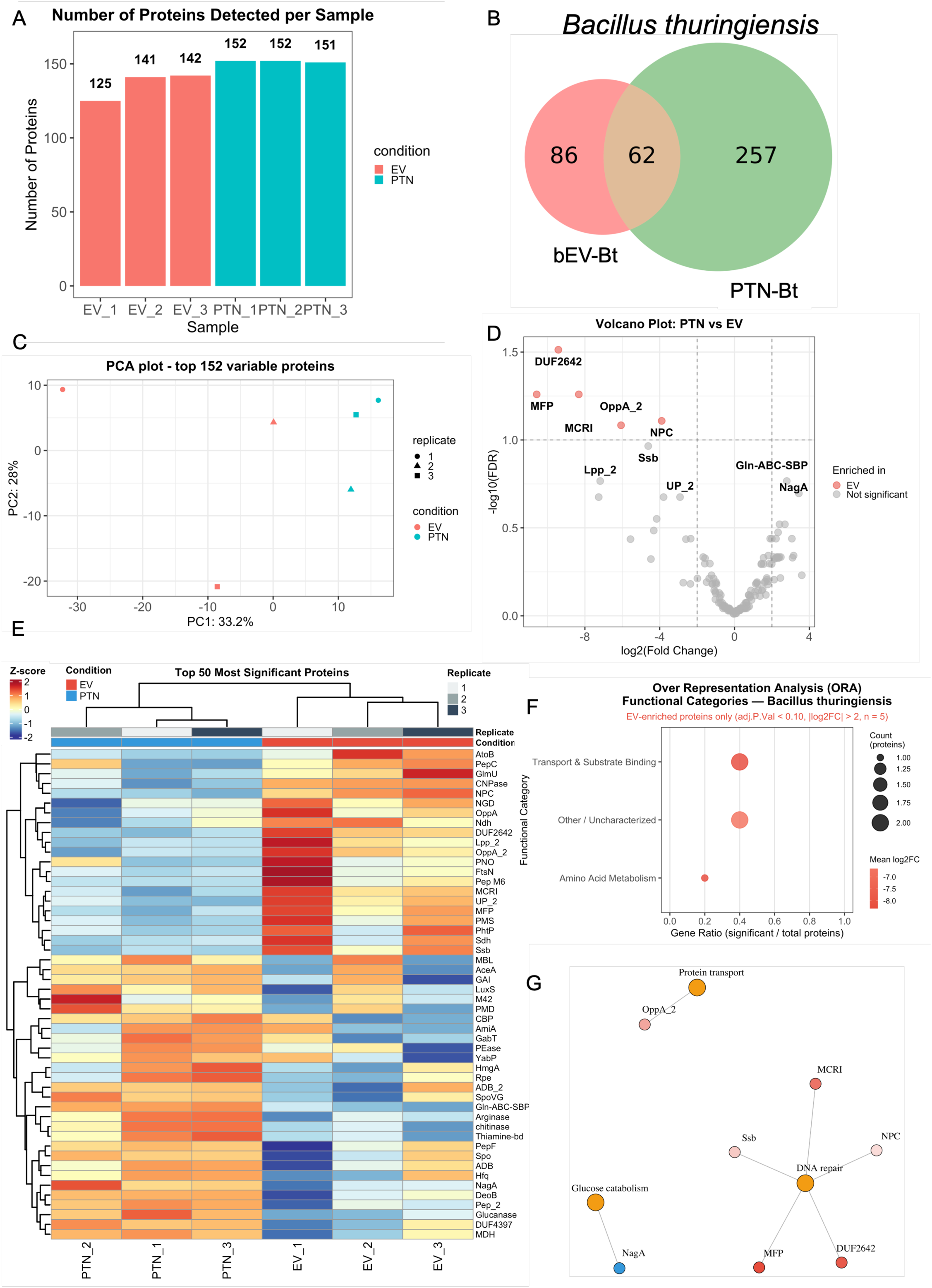
Quantitative proteomics of extracellular vesicles (bEVs) and soluble protein fractions from *Bacillus thuringiensis*. (A) Number of proteins detected in each biological replicate of *B. thuringiensis* bEV samples (EV_1–EV_3) and soluble protein fractions (PTN_1–PTN_3). (B) Venn diagram of proteins detected exclusively in, or shared between, the bEV-enriched (bEV-Bt) and soluble protein (PTN-Bt) fractions. (C) Principal component analysis (PCA) of the top 152 variable proteins across all *B. thuringiensis* samples, coloured by condition and shaped by replicate. (D) Volcano plot of log₂(fold-change) versus −log₁₀(adjusted p-value) for differential protein abundance between the PTN and EV fractions. Labeled proteins are the most differentially abundant; enrichment reaching significance was observed only in the EV fraction. (E) Hierarchically clustered heatmap of the 50 most significant proteins across all replicates, organized by condition (PTN and EV), with z-score normalization indicated by the colour scale. (F) Gene Ontology over-representation analysis (ORA) of *B. thuringiensis* bEV-enriched proteins recovered under the exploratory threshold (adjusted p < 0.10, |log₂FC| > 2; n = 5). Bubble size, proteins per category; colour, mean log₂(fold-change). (G) Protein–functional-category network of the most differentially abundant *B. thuringiensis* proteins, organized by functional module; node size proportional to −log₁₀(p-value), edges representing functional relationships.

Presence/absence analysis detected 86 proteins only in the bEV fraction, 257 only in the soluble protein fraction, and 62 shared between fractions (**Figure 4B**). Because data-dependent acquisition can lead to stochastic non-detection of low-abundance proteins, these presence/absence categories were treated as provisional, and the interpretation of enrichment was based primarily on formal differential abundance analysis.

Quality-control analyses supported downstream statistical comparison. Missing-value counts were low across replicates (**Supplementary Figure 7A**), variance-stabilizing normalization balanced intensity distributions between bEV and soluble protein fractions (**Supplementary Figure 7B**), and comparison of imputation strategies supported the use of MinProb imputation for downstream analysis (**Supplementary Figure 7C**). Principal component analysis separated bEV and soluble protein samples along PC1, which explained 33.2% of the variance, although within-group replicate dispersion was observed (**Figure 4C**; **Supplementary Figure 7D,E**). The complete *B. thuringiensis* quantitative dataset and differential-abundance statistics are provided in **Supplementary Tables 1 and 2**. Differential-abundance analysis showed limited proteomic separation between *B. thuringiensis* bEV and soluble protein fractions. Of 152 quantified proteins, only one protein, DUF2642, was significantly enriched in the bEV fraction at FDR < 0.05 and |log₂FC| > 1 (log₂FC = −9.42, adjusted p = 0.0307; **Figure 4D**; **Supplementary Table 2**). Using an exploratory relaxed threshold of FDR < 0.10 identified four additional bEV-enriched candidates: MCRI, MFP, NPC, and OppA_2. Several proteins showed large negative log₂FC values but did not meet the FDR threshold, including Ssb and Lpp_2. No proteins were significantly enriched in the soluble protein fraction at FDR < 0.05 and |log₂FC| > 1. Because *B. thuringiensis* var. *kurstaki* is associated with Cry toxin production, the proteomics dataset was further examined for toxin-related annotations. A crystalline entomocidal protoxin annotated as Cry1Ab was detected in the total proteomics report with 13 peptides. This protein showed higher mean normalized abundance in the soluble protein fraction than in the bEV fraction, with the highest mean condition assigned to PTN and an approximately 62.9-fold difference in the total report (**Supplementary Table 1**). In the formal differential-abundance table, the corresponding toxin-related feature did not meet the significance threshold after FDR correction (**Supplementary Table 2**). No entries annotated as Vip, Cyt, hemolysin, cytolysin, RTX toxin, or adenylate-cyclase toxin were detected in the *B. thuringiensis* proteomics tables.

Additional extracellular or barrier-associated protein classes were detected in the *B. thuringiensis* dataset. The total proteomics report included chitin-associated proteins, including two chitinases and two chitin-binding proteins, all with the highest mean condition assigned to the soluble protein fraction (**Supplementary Table 1**). Peptidases, aminopeptidases, hydrolases, and glucanase-related proteins were also detected, including Peptidase M28, Xaa-Pro aminopeptidase, Dipeptidase PepV, Aminopeptidase, Peptidase M42, Peptidase M24, Oligopeptidase F, neutral metalloproteinase, amidohydrolases, metal-dependent hydrolases, and glucanase entries (**Supplementary Table 1**). In the formal DEP analysis, these proteins did not meet the prespecified FDR < 0.05 and |log₂FC| > 1 threshold (**Supplementary Table 2**).

Hierarchical clustering of the 50 proteins with the lowest unadjusted p-values showed partial separation between bEV and soluble protein fractions, consistent with the limited number of statistically significant differentially abundant proteins (**Figure 4E**). Gene Ontology over- representation analysis was performed using the five bEV-enriched proteins recovered under the exploratory FDR < 0.10 threshold and identified categories related to transport and substrate binding, amino acid metabolism, and other/uncharacterized proteins (**Figure 4F**). The corresponding protein–functional-category network showed a sparse structure centered on the small set of exploratory bEV-enriched candidates (**Figure 4G**). Overall, the *B. thuringiensis* proteomic data showed limited selective remodeling between bEV and soluble protein fractions, with DUF2642 as the only protein significantly enriched in bEVs under the prespecified statistical criteria.

### *C. subtsugae* bEVs show selective proteomic remodeling and enrichment of candidate host-interaction proteins

Label-free quantitative proteomics was performed on *C. subtsugae* bEV-enriched and soluble protein fractions using three independent biological replicates per fraction. bEV preparations yielded 362, 363, and 359 protein identifications, whereas soluble protein fractions yielded 370, 362, and 367 identifications (**Figure 5A**). Presence/absence analysis detected 244 proteins only in the bEV fraction, 282 only in the soluble protein fraction, and 185 shared between fractions (**Figure 5B**). As for *B. thuringiensis*, these presence/absence categories were treated as provisional because data-dependent acquisition can lead to stochastic non-detection of low-abundance proteins; therefore, enrichment interpretation was based primarily on formal differential-abundance analysis.

**Figure 5.**
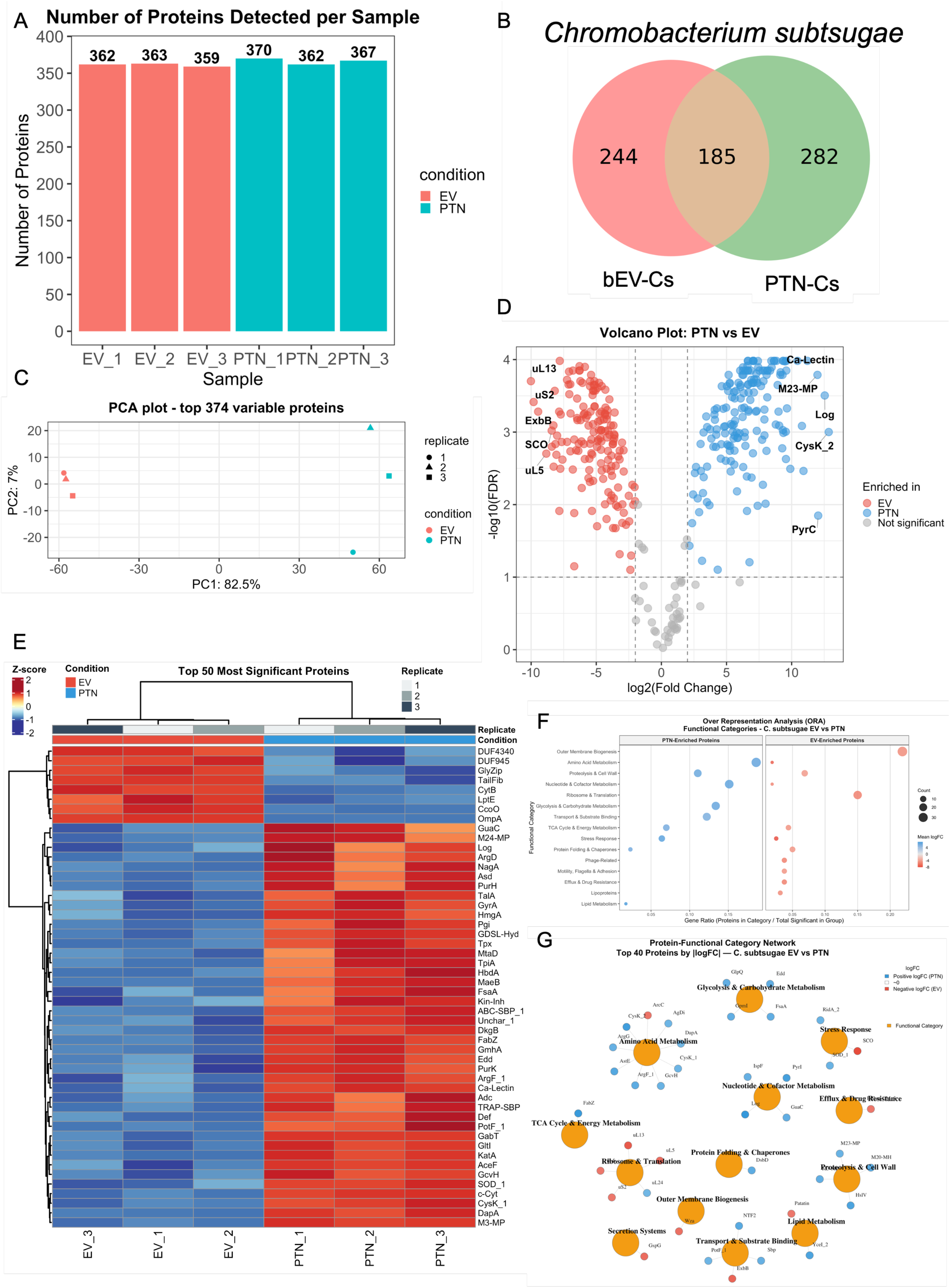
Quantitative proteomics of extracellular vesicles (bEVs) and soluble protein fractions from *C. subtsugae*. (A) Number of proteins detected in each biological replicate of *C. subtsugae* bEV samples (EV_1–EV_3) and soluble protein fractions (PTN_1–PTN_3). (B) Venn diagram of proteins detected exclusively in, or shared between, the bEV-enriched (bEV-Cs) and soluble protein (PTN-Cs) fractions. (C) Principal component analysis (PCA) of the top 374 variable proteins across all *C. subtsugae* samples, coloured by condition and shaped by replicate. (D) Volcano plot of log₂(fold-change) versus −log₁₀(adjusted p-value) for differential protein abundance between the PTN and EV fractions. Labeled proteins are the most differentially abundant in each fraction (red, EV-enriched; blue, PTN-enriched). (E) Hierarchically clustered heatmap of the 50 most significant proteins across all replicates, organized by condition (PTN and EV), with z-score normalization indicated by the colour scale. (F) Gene Ontology over-representation analysis (ORA) of differentially abundant *C. subtsugae* proteins, shown separately for PTN-enriched and EV-enriched functional categories. Bubble size, proteins per category; colour, mean log₂(fold-change). (G) Protein–functional-category network of the top 40 proteins by |log₂(fold-change)|, coloured by fraction of enrichment (red, EV-enriched; blue, PTN-enriched); node size proportional to −log₁₀(p-value), edges linking proteins to functional modules.

Quality-control analyses supported robust downstream comparison. Missing-value counts were low across replicates (**Supplementary Figure 8A**), variance-stabilizing normalization balanced intensity distributions between bEV and soluble protein fractions (**Supplementary Figure 8B**), and comparison of imputation strategies supported the use of MinProb imputation for downstream analysis (**Supplementary Figure 8C**). Principal component analysis revealed pronounced separation between bEV and soluble protein fractions along PC1, which explained 82.5% of the variance, with complete segregation of all biological replicates by fraction (**Figure 5C**; **Supplementary Figure 8D,E**). This separation showed that the *C. subtsugae* bEV-enriched fraction had a proteomic profile that was clearly distinct from the soluble protein-enriched fraction. The complete quantitative dataset and differential-abundance statistics are provided in **Supplementary Tables 3 and 4**.

Differential-abundance analysis confirmed extensive compositional remodeling between fractions. Of 374 quantified proteins, 334 were significantly differentially abundant at FDR < 0.05 and |log₂FC| > 1, comprising 162 proteins enriched in the bEV fraction and 172 enriched in the soluble protein fraction (**Figure 5D**; **Supplementary Table 4**). Negative log₂FC values indicated enrichment in the bEV fraction, whereas positive log₂FC values indicated enrichment in the soluble protein fraction. Hierarchical clustering of the most significantly differentially abundant proteins reproduced this fraction-dependent separation across all biological replicates (**Figure 5E**).

The bEV-enriched proteome contained multiple proteins consistent with an outer-membrane vesicle-associated profile. These included OmpA-family proteins, OmpH, LptE, ExbB, Wza, TolC-family proteins, outer-membrane efflux components, porins, and lipoproteins. Several proteins related to envelope biogenesis and membrane-associated transport were also enriched in the bEV fraction, including outer-membrane assembly factors and lipoprotein-associated proteins. In addition, the bEV fraction contained secretion-associated proteins, including the type II secretion system protein GspG and the type VI secretion system lipoprotein TssJ, and was enriched in several proteolytic or peptidase-related proteins, including M48 family metalloproteases, DegP-like periplasmic serine endoproteases, S41 and S49 family peptidases, LdcA, and MmpCys. A patatin-like phospholipase family protein was also significantly enriched in the bEV fraction (**Supplementary Table 4**). No chitinase or chitin-binding protein annotations were detected among the *C. subtsugae* proteomics entries. A translation-associated module was likewise observed among the bEV-enriched proteins, with 24 of 30 identified ribosomal subunits showing higher abundance in the bEV fraction.

The soluble protein-enriched fraction displayed a different functional profile. Proteins enriched in this fraction included enzymes associated with central carbon metabolism, glycolysis and carbohydrate metabolism, amino acid biosynthesis and catabolism, nucleotide metabolism, antioxidant response, and proteolysis. Examples included FabZ, GpmI, FsaA, Edd, CysK, ArgF, ArgG, GuaC, PyrI, KatA, SOD, HslV, and several additional metabolic enzymes and uncharacterized proteins.

Classical secreted toxin annotations, including pore-forming cytolysins, RTX/hemolysins, and adenylate-cyclase toxins, were not detected among the differentially abundant proteins in either fraction. Consistent with the protein-level observations, functional enrichment analysis highlighted outer-membrane biogenesis, transport and substrate binding, secretion systems, and translation among bEV-associated categories, whereas central metabolism was preferentially associated with the soluble protein fraction (**Figure 5F**). The corresponding protein–functional-category network resolved these relationships into discrete functional modules, with envelope-, secretion-, and translation-associated proteins clustering separately from metabolic enzymes (**Figure 5G**). Overall, these data show that *C. subtsugae* bEVs represent a selectively remodeled vesicular compartment enriched in envelope-associated, secretion-associated, proteolytic, and membrane-active proteins.

### bEV and soluble protein fractions display species-specific insecticidal activity against Euschistus heros

The insecticidal activity of bacterial bEVs and soluble protein fractions was evaluated using a 168 h Kaplan–Meier survival assay with second-instar *E. heros* nymphs. Each treatment was compared against the diet-only negative control (NC), and Grandevo®, a commercial *C. subtsugae*-based biopesticide, was included as a positive control (PC). Survival curves for each treatment are shown in **Figure 6A–F**.

**Figure 6.**
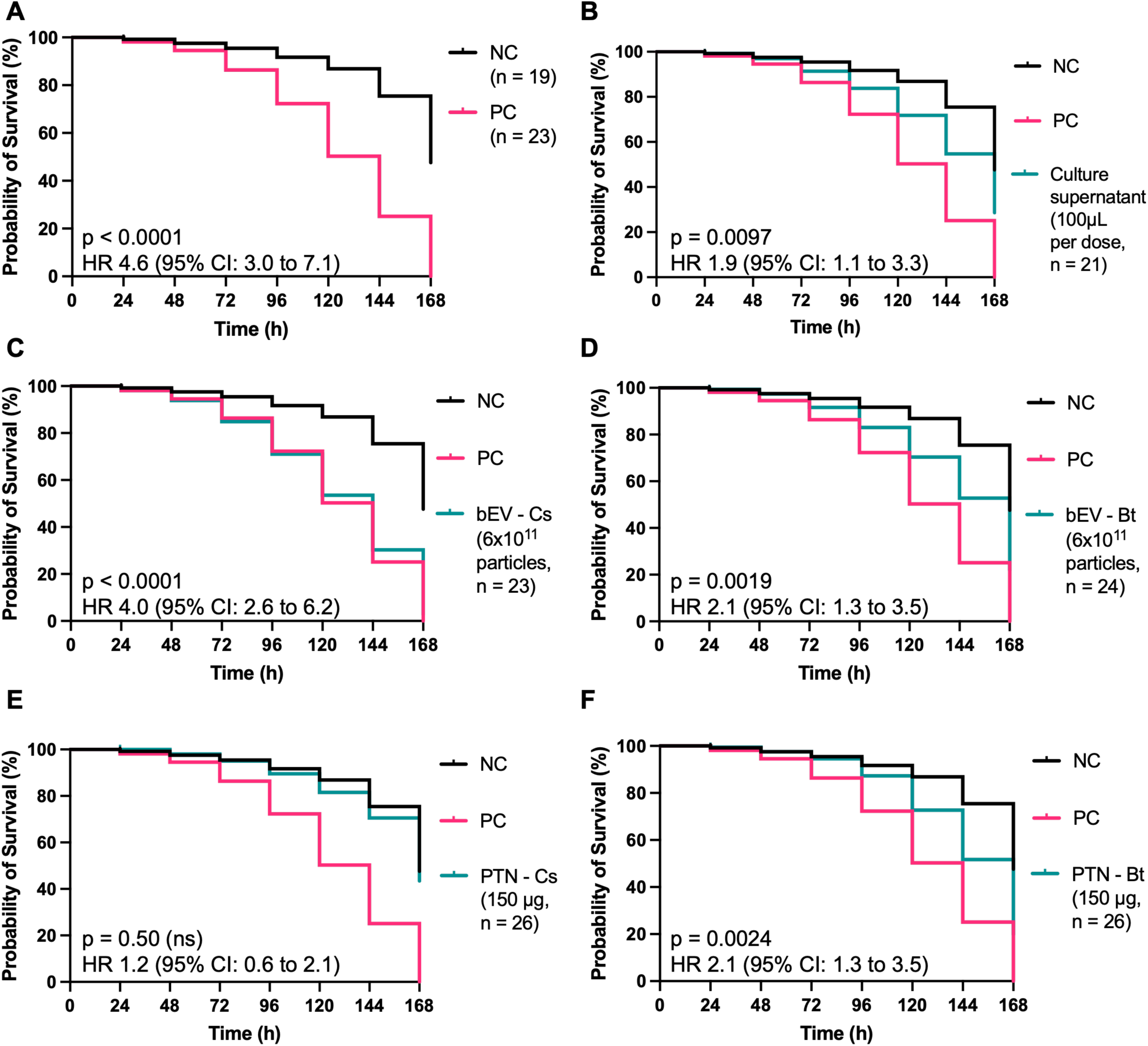
Insecticidal activity of bacterial extracellular vesicles (bEVs) and soluble protein fractions against *E. heros*. (A) Kaplan–Meier survival curve comparing the negative control (NC; sterile diet) with the positive control (PC; Grandevo®, 350 µg of formulated product per dose). (B) Kaplan–Meier survival curve comparing NC and PC with whole *C. subtsugae* culture supernatant (100 µL per dose). (C) Kaplan–Meier survival curve comparing NC and PC with isolated *C. subtsugae* bEVs (6 × 10¹¹ particles, ∼284 µg protein per dose). (D) Kaplan–Meier survival curve comparing NC and PC with isolated *B. thuringiensis* bEVs (6 × 10¹¹ particles, ∼136 µg protein per dose). (E) Kaplan–Meier survival curve comparing NC and PC with the *C. subtsugae* soluble protein fraction (PTN-Cs; 150 µg per dose). (F) Kaplan–Meier survival curve comparing NC and PC with the *B. thuringiensis* soluble protein fraction (PTN-Bt; 150 µg per dose). All curves show *E. heros* second-instar nymph survival monitored over 168 h; n values indicate the number of nymphs alive and feeding at treatment onset (t = 0) after the 48-hour acclimatization period. Log-rank (Mantel–Cox) p-values and hazard ratios (Mantel–Haenszel method) with 95% confidence intervals, each relative to NC, are indicated on each panel.

The positive control induced a strong reduction in nymph survival, approaching near-complete mortality by 168 h (**Figure 6A**). Log-rank analysis showed a significant difference between PC and NC, with HR = 4.6, 95% CI: 3.0–7.1, and p < 0.0001. Crude *C. subtsugae* culture supernatant, containing both vesicular and soluble components at their native ratio, produced an intermediate reduction in survival (**Figure 6B**). This effect was significant relative to NC, with HR = 1.9, 95% CI: 1.1–3.3, and p = 0.0097.

Purified *C. subtsugae* bEVs, administered at 6 × 10¹¹ vesicles and approximately 284 µg protein per dose, markedly reduced *E. heros* nymph survival (**Figure 6C**). The strongest decline occurred between 72 and 120 h, with minimal survival remaining by 168 h. This effect was significant relative to NC, with HR = 4.0, 95% CI: 2.6–6.2, and p < 0.0001. The magnitude of this effect was close to that observed for the Grandevo® positive control. Purified *B. thuringiensis* bEVs, administered at 6 × 10¹¹ vesicles and approximately 136 µg protein per dose, also reduced nymph survival relative to NC (**Figure 6D**). The strongest reduction occurred between 96 and 144 h. Log-rank analysis showed a significant difference relative to NC, with HR = 2.1, 95% CI: 1.3–3.5, and p = 0.0019. The effect magnitude was lower than that observed for *C. subtsugae* bEVs.

The *C. subtsugae* soluble protein fraction, administered at 150 µg protein per dose, did not significantly alter nymph survival relative to NC (**Figure 6E**). The estimated HR was 1.2, with a 95% CI of 0.6–2.1 and p = 0.50. In contrast, the *B. thuringiensis* soluble protein fraction, also administered at 150 µg protein per dose, significantly reduced nymph survival (**Figure 6F**). The effect was similar to that observed for *B. thuringiensis* bEVs, with HR = 2.1, 95% CI: 1.3–3.5, and p = 0.0024.

Overall, the survival assay showed distinct activity profiles between the two bacterial species. For *C. subtsugae*, insecticidal activity was strongly associated with the bEV- enriched fraction, whereas the soluble protein fraction was inactive under the conditions tested. For *B. thuringiensis*, both bEV and soluble protein fractions produced similar moderate reductions in nymph survival.

## Discussion

Bacterial extracellular vesicles have emerged as important mediators of microbial communication, environmental adaptation, and delivery of bioactive molecules. In bacteria, EVs can transport proteins, lipids, nucleic acids, toxins, enzymes, and secondary metabolites, allowing microorganisms to influence neighboring cells and host organisms without requiring direct cell-to-cell contact (Kim et al., 2015)(Sartorio et al., 2021)(Toyofuku et al., 2023). Although EVs have been extensively investigated in mammalian systems and increasingly in plant–microbe interactions, their role in agricultural pest control remains poorly understood. Recent studies have shown that EVs from plant-associated bacteria can modulate interkingdom communication, promote plant growth, and influence plant resistance to pathogens (Zhou et al., 2022)(Zannis-Peyrot et al., 2025). However, the possibility that purified bEVs from entomopathogenic bacteria can function as bioinsecticidal particles remains largely unexplored.

Here, we provide proof-of-concept evidence that enriched bEVs from entomopathogenic bacteria can act as non-replicative bioinsecticidal particles against the soybean pest *Euschistus heros*. This extends the early observation that purified OMVs from *Xenorhabdus nematophilus* possess intrinsic insecticidal activity (Khandelwal and Banerjee-Bhatnagar, 2003) to two commercially relevant bacterial species, *Chromobacterium subtsugae* and *Bacillus thuringiensis*. Importantly, our data indicate that bEV-associated insecticidal activity is species-specific: *C. subtsugae* activity was strongly associated with the bEV-enriched fraction, whereas *B. thuringiensis* activity was distributed similarly between bEV and soluble protein fractions. These contrasting profiles suggest that bEV-based bioactivity depends on how each bacterium distributes bioactive metabolites and proteins across vesicular and non-vesicular extracellular compartments.

The strongest evidence for vesicle-associated bioinsecticidal activity was observed for *C. subtsugae*. Purified *C. subtsugae* bEVs markedly reduced *E. heros* nymph survival, with an effect approaching that of the commercial *C. subtsugae*-based biopesticide Grandevo®, whereas the corresponding soluble protein fraction, derived from the same fermentation batches and fractionation workflow, was inactive under the same assay conditions. Although the two fractions were not dose-matched by total protein (approximately 284 µg for the bEV- enriched fraction versus 150 µg for the soluble protein fraction), this difference does not account for the contrasting outcomes: the soluble fraction produced no detectable effect, with a hazard ratio confidence interval spanning 1.0, indicating an absence of activity rather than a reduced-magnitude response. This result suggests that the insecticidal activity of *C. subtsugae* is preferentially associated with the vesicular compartment rather than with freely soluble extracellular proteins.

This finding is consistent with prior work showing that *Chromobacterium* species can use OMVs to package and export hydrophobic bioactive compounds, including violacein (Choi et al., 2020). In *C. violaceum*, violacein secretion and OMV biogenesis are influenced by quorum-sensing-regulated pathways, linking pigment production, vesiculation, and microbial interaction phenotypes (Batista et al., 2020)(Devescovi et al., 2017).

Violacein is a strong candidate contributor to the activity observed here. This indole-derived hydrophobic metabolite has been associated with antimicrobial, antiparasitic, cytotoxic, and insecticidal activities (Martin et al., 2007)(Chauhan et al., 2025)(Scaramussa et al., 2025). Previous studies showed that membrane vesicles can improve the delivery of poorly soluble secondary metabolites, including violacein, to target cells or organisms (Kowalska et al., 2024) (Jorge et al., 2025). Similar principles have been described for other hydrophobic bacterial metabolites, such as phenazines in *Pseudomonas*, where vesiculation can facilitate transport, stabilization, or delivery of poorly soluble compounds (Bitzenhofer et al., 2024).

In our study, MALDI-TOF MS detected a violacein-associated ion selectively in the *C. subtsugae* bEV fraction, but not in the corresponding soluble protein fraction. *B. thuringiensis* preparations, which served as negative specificity controls given that this species lacks the violacein biosynthetic pathway, likewise showed no such ion, confirming that the signal was not attributable to matrix or medium-derived background. This supports a model in which OMVs provide a lipid-based compartment for concentrating and delivering violacein-like hydrophobic cargo.

However, the activity of *C. subtsugae* bEVs is unlikely to be explained by violacein alone. Instead, our data support a multi-component cargo model in which vesicle-associated violacein, additional hydrophobic metabolites, and EV-enriched proteins may act together to produce insecticidal activity. This interpretation is supported by the strong proteomic separation between *C. subtsugae* bEV and soluble protein fractions, with PC1 explaining 82.5% of the variance. Differential-abundance analysis further showed extensive compositional remodeling, with 334 of 374 quantified proteins significantly differentially abundant between fractions, including 162 proteins enriched in bEVs and 172 enriched in the soluble protein fraction. Thus, the *C. subtsugae* bEV fraction does not appear to be a simple reflection of the soluble secretome, but rather a distinct vesicle-associated compartment.

The *C. subtsugae* bEV-enriched proteome contained several classes of proteins that may be relevant to host interaction. Outer-membrane and envelope-associated proteins, including OmpA-family proteins, LptE, ExbB, Wza, TolC-family proteins, outer-membrane efflux components, porins, and lipoproteins, support the identity of this fraction as an OMV-enriched preparation. This agrees with the expected composition of Gram-negative OMVs, which commonly carry outer-membrane, periplasmic, and envelope-associated proteins (Sartorio et al., 2021)(Orench-Rivera and Kuehn, 2021). The enrichment of secretion-associated proteins such as GspG and TssJ also suggests that the *C. subtsugae* bEV fraction contains host-interaction-associated components, consistent with the broader role of bacterial EVs in delivering virulence and interaction factors (Kim et al., 2015)(Toyofuku et al., 2023).

Notably, the *C. subtsugae* bEV fraction was enriched in several proteolytic or membrane-active proteins, including M48 family metalloproteases, DegP-like serine endoproteases, S41 and S49 family peptidases, LdcA, MmpCys, and a patatin-like phospholipase. These proteins are not classical insecticidal toxins, but they are biologically plausible accessory effectors. In other bacterial systems, EV-associated proteases and membrane-active enzymes can disrupt host barriers and enhance delivery of bacterial cargo. For example, *Pseudomonas aeruginosa* OMVs carry LasB elastase, a metalloprotease associated with tissue damage and disruption of epithelial barriers (Galdino et al., 2019)(Zhou et al., 2025). *Helicobacter pylori* OMVs can carry HtrA, a serine protease that contributes to disruption of epithelial junctions (Zawilak-Pawlik et al., 2019). Similarly, *Vibrio cholerae* OMV-associated HapA has been linked to disruption of tight and adherens junctions (Baryalai et al., 2025). In Gram-positive bacteria, *Bacillus cereus* EVs can deliver biologically active multicomponent enterotoxins to host cells (Buchacher et al., 2023). By analogy, the proteases, peptidases, and phospholipase enriched in *C. subtsugae* bEVs may contribute to interaction with insect tissues, membrane perturbation, barrier disruption, or enhanced access of vesicle-associated metabolites such as violacein. This remains a biologically plausible mechanistic hypothesis that warrants future functional testing.

The enrichment of ribosomal proteins in the *C. subtsugae* bEV-enriched fraction requires careful interpretation. Ribosomal and other cytoplasmic proteins are frequently flagged as indicators of non-vesicular carryover in EV preparations. This consideration is particularly relevant here because supernatants were collected at stationary phase, when cell death and lysis are expected to increase. Cell disruption can release cytoplasmic content into the culture medium and, as noted in MISEV2023 (Welsh et al., 2024), can also generate particles that resemble native EVs in size and biophysical properties and that co-purify through the same isolation workflow. Consequently, lysis-derived material cannot be excluded as a contributor to the observed proteome (Turnbull et al., 2016).

At the same time, the ribosomal signature occurred within a broader EV-associated proteomic profile that was reproducibly separated from the soluble protein fraction by PCA, differential-abundance analysis, and functional enrichment, and that was dominated by outer-membrane, envelope, transport, and secretion-associated proteins consistent with bona fide OMV composition. We note, however, that reproducible separation alone does not discriminate between selective vesicular packaging and systematic co-isolation of lysis-derived particles, since both would be expected to partition consistently across replicates. These observations are therefore compatible with more than one interpretation: selective incorporation of cytoplasm-derived material during vesicle biogenesis, lysis-associated vesiculation such as explosive cell lysis, or co-isolation of non-vesicular cytoplasmic material. Distinguishing among these possibilities will require density-gradient fractionation, protease-protection assays, and comparison of vesicle production across growth phases. In contrast to *C. subtsugae*, *B. thuringiensis* showed limited selective remodeling between bEV and soluble protein fractions. Only DUF2642 was significantly enriched in the bEV fraction under the prespecified statistical threshold. DUF-annotated proteins remain common in bacterial proteomes, comprising more than 20% of predicted bacterial proteins, and some have subsequently been shown to perform specific, biologically important functions once structurally or functionally characterized (Patel et al., 2025), the biological role of DUF2642 in this context therefore remains an open question for future investigation. This limited proteomic separation is consistent with the bioassay results, in which *B. thuringiensis* bEVs and soluble protein fractions produced similar moderate reductions in *E. heros* survival.

Thus, under the conditions tested, *B. thuringiensis* insecticidal activity against *E. heros* was not selectively concentrated in the bEV fraction. This result is particularly relevant because the *B. thuringiensis* strain used here belongs to var. *kurstaki*, which is classically associated with Cry1Ab production and activity against lepidopteran larvae (Bravo et al., 2007). Conventional Cry toxicity depends on ingestion of crystalline protoxins, solubilization and proteolytic activation in the insect gut, receptor binding in the midgut epithelium, pore formation, and epithelial cell lysis (Kumar et al., 2021). Hemipteran sap-feeding insects such as *E. heros* are generally poorly targeted by conventional Bt Cry toxins because their feeding behavior, gut physiology, toxin-processing environment, and receptor interactions differ from those of chewing lepidopteran larvae (Li et al., 2011)(Chougule and Bonning, 2012)(Rausch et al., 2016). This biological mismatch helps explain why Cry1Ab-based *B. thuringiensis* var. *kurstaki* mechanisms are not expected to be highly effective against *E. heros*.

In agreement with this expectation, Cry1Ab was detected in the *B. thuringiensis* proteomics dataset but was more abundant in the soluble protein fraction than in the bEV fraction. Moreover, no Vip, Cyt, RTX, hemolysin, cytolysin, or adenylate-cyclase toxin annotations were detected in the *B. thuringiensis* proteomics tables. Although Vip3Aa has previously been reported to associate with *B. thuringiensis* membrane vesicles (Zhang et al., 2022), our data do not support broad enrichment of classical Bt toxins in the bEV fraction of the strain and growth condition tested here. Instead, the moderate activity observed for *B. thuringiensis* may reflect non-classical extracellular factors distributed across vesicular and soluble compartments. The detection of chitinases, chitin-binding proteins, peptidases, aminopeptidases, hydrolases, and glucanase-related proteins mainly in the soluble protein-enriched fraction supports this interpretation.

Under the conditions tested, the two species showed different distributions of insecticidal activity between the early bEV-enriched and later soluble-protein-enriched SEC fractions. In *C. subtsugae*, insecticidal activity was strongly associated with the bEV-enriched fraction, which combined selective detection of a violacein-associated signal with a remodeled proteome enriched in envelope-associated, secretion-associated, proteolytic, and membrane-active proteins.

In *B. thuringiensis*, activity was shared between bEV and soluble protein fractions, while Cry1Ab and several barrier-associated enzymes were detected mainly in the soluble protein-enriched fraction. This contrast reinforces the idea that bEVs do not represent a universal delivery mechanism for all bacterial insecticidal factors; rather, their bioactivity depends on species-specific cargo partitioning and the chemical nature of the active molecules.

From a biotechnological perspective, these findings support bEVs as a promising platform for developing non-replicative bioinsecticides. bEVs combine two attractive properties: they are cell-free particles, and they can carry complex bioactive cargo derived from the producing bacterium (Kim et al., 2015)(Sartorio et al., 2021)(Toyofuku et al., 2023). This distinguishes them from single-molecule bioinsecticides and may be particularly relevant in pest-management contexts where multi-component activities could reduce the likelihood of rapid resistance evolution. EV-based agricultural applications are still emerging, but recent studies support their involvement in microbial communication, plant–microbe interactions, plant protection, and interkingdom delivery processes (Xie et al., 2022)(González and Falcón-Pérez, 2025)(Remans et al., 2025). More broadly, current EV frameworks also recognize extracellular vesicles as stable and biologically active delivery systems across microbial, plant, and animal systems (Toyofuku et al., 2023)(Welsh et al., 2024). In this context, violacein-associated *C. subtsugae* OMVs provide a compelling proof-of-concept for natural bacterial nanocarriers in insect pest control.

The proof-of-concept demonstrated here is particularly relevant for *E. heros*, a major hemipteran soybean pest for which sustainable biological control options remain limited (Ferreira Agüero et al., 2023)(Ferreira Agüero et al., 2023)(Saldanha et al., 2024). This species has evolved resistance-associated mechanisms against conventional insecticides, including target-site mutations affecting GABA-gated chloride channels (Cuenca et al., 2025), and is not expected to be highly susceptible to classical Cry1Ab-based *Bt* activity because hemipteran feeding behavior, gut physiology, toxin processing, and receptor interactions differ from those of lepidopteran larvae (Li et al., 2011)(Chougule and Bonning, 2012)(Rausch et al., 2016). The activity observed with *C. subtsugae* bEVs suggests that vesicle-associated metabolites and proteins may bypass some of the biological constraints that limit conventional Bt toxin efficacy in hemipterans. These findings do not establish bEVs as ready-to-use field bioinsecticides, but they establish a foundation for future development of vesicle-based formulations and for mechanistic studies of bEV-mediated insecticidal delivery.

## Limitations of the study

While this study provides, to our knowledge, the first evidence for bEV-associated bioinsecticidal activity against *E. heros*, three limitations should be considered.

First, the mechanisms by which bEVs are internalized and exert lethal effects in *E. heros* remain unknown. Here, bEVs were delivered through an artificial diet, whereas under field conditions *E. heros* feeds on plant sap using piercing-sucking mouthparts. Therefore, it remains unclear whether bEVs would reach the insect through foliar contact, plant tissue colonization, or other exposure routes. For *C. subtsugae*, violacein is a leading candidate effector, but direct evidence of bEV uptake and violacein release in insect tissues is still lacking. For *B. thuringiensis*, the active component shared between vesicular and soluble fractions remains unresolved.

Second, the bioassays were performed using a single dose and only second-instar *E. heros* nymphs. Dose-response assays will be required to determine EC50 or LC50 values and compare bEV potency with existing chemical and biological insecticides. Additional tests across developmental stages, pest species, and non-target organisms, including beneficial insects and pollinators, will also be necessary to define the activity spectrum and safety profile of bEV-based bioinsecticides.

Third, translation from controlled bioreactor production to field-applicable formulations remains challenging. Future work must address scalable bEV production, formulation stability under environmental stressors such as UV radiation, temperature variation, and pH fluctuation, and standardized crop-application protocols before bEV-based bioinsecticides can be considered for practical pest-management programs.

All three limitations are currently being addressed in ongoing work by our group.

## Conclusion

This study provides proof-of-concept evidence that bacterial extracellular vesicles can function as non-replicative bioinsecticidal particles against the soybean pest *Euschistus heros*. *C. subtsugae* bEVs showed strong insecticidal activity and combined selective detection of a violacein-associated signal with a remodeled vesicular proteome enriched in envelope-associated, secretion-associated, proteolytic, and membrane-active proteins. These findings support a multi-component cargo model in which vesicle-associated metabolites and proteins may jointly contribute to insecticidal activity. In contrast, *B. thuringiensis* bEV and soluble protein fractions showed similar moderate activity, consistent with limited selective remodeling of the bEV proteome and with Cry1Ab being detected mainly in the soluble fraction. Overall, these results establish bEVs as promising natural nanocarriers for bioinsecticidal cargo and support their further development as sustainable, cell-free platforms for pest management.

## Acknowledgments

This work was supported by the Fundação de Apoio à Pesquisa do Distrito Federal (FAP- DF; Chamada Pública FAPDF No. 01/2024 – PDPG Rede de Pesquisa e Desenvolvimento da Região Centro-Oeste; to MSSF), the Conselho Nacional de Desenvolvimento Científico e Tecnológico (CNPq; Chamada CNPq/MCTI/FNDCT No. 21/2024 – Programa Conhecimento Brasil: Atração e Fixação de Talentos; to GPOJ), and the Financiadora de Estudos e Projetos (FINEP; to SNB). Graduate and undergraduate fellowships were provided by the Coordenação de Aperfeiçoamento de Pessoal de Nível Superior (CAPES, Brazil). Institutional support and research facilities were provided by the Universidade Católica de Brasília. Transmission electron microscopy was performed at the Laboratório de Microscopia e Microanálise of the Universidade de Brasília.

## Data availability statement

The raw LC-MS/MS proteomics data generated during this study have been deposited in the MassIVE repository under accession number MSV000102695 and are available through the ProteomeXchange Consortium under accession number PXD082005. Processed proteomics results, including protein identification, quantification, and differential-abundance analyses, are provided in the Supplementary Tables. Additional supporting data, including MALDI-TOF mass spectrometry analyses, bEV characterization data, bioassay protocols, and statistical analyses, are included in the main article and Supplementary Information. Other data relevant to this study are available from the corresponding authors upon reasonable request.

## Declaration of generative AI in scientific writing

During the preparation of this work, GPOJ used Claude Pro (Anthropic) in order to improve language and readability of the manuscript. After using this tool, the authors reviewed and edited the content as needed and take full responsibility for the content of the publication.

## References

Baryalai, Palwasha et al. Hemagglutinin Protease HapA Associated With *Vibrio cholerae* Outer Membrane Vesicles (OMVs) Disrupts Tight and Adherens Junctions. Journal of Extracellular Vesicles, v. 14, n. 5, 25 maio 2025.

Batista, Juliana H. et al. Interplay between two quorum sensing-regulated pathways, violacein biosynthesis and VacJ /Yrb, dictates outer membrane vesicle biogenesis in *Chromobacterium violaceum*. Environmental Microbiology, v. 22, n. 6, p. 2432–2442, 5 jun. 2020.

Bitzenhofer, Nora Lisa et al. Exploring engineered vesiculation by *Pseudomonas putida* KT2440 for natural product biosynthesis. Microbial Biotechnology, v. 17, n. 1, 12 jan. 2024.

Bravo, Alejandra; Gill, Sarjeet S.; Soberón, Mario. Mode of action of Bacillus thuringiensis Cry and Cyt toxins and their potential for insect control. Toxicon, v. 49, n. 4, p. 423–435, mar. 2007.

Briaud, Paul; Carroll, Ronan K. Extracellular Vesicle Biogenesis and Functions in Gram-Positive Bacteria. Infection and Immunity, v. 88, n. 12, 16 nov. 2020.

Buchacher, Tanja et al. Bacillus cereus extracellular vesicles act as shuttles for biologically active multicomponent enterotoxins. Cell Communication and Signaling, v. 21, n. 1, p. 112, 15 maio 2023.

Castro, Marcelo T. de et al. Susceptibility of Euschistus heros (Fabricius, 1798) (Hemiptera: Pentatomidae) nymphs to Bacillus spp. strains. Entomological Communications, v. 7, p. ec07028, 2 out. 2025.

Chauhan, Abhishek et al. Mechanistic Insight of Pharmacological Aspects of Violacein: Recent Trends and Advancements. Journal of Biochemical and Molecular Toxicology, v. 39, n. 2, 26 fev. 2025.

Choi, Seong Yeol et al. *Chromobacterium violaceum* delivers violacein, a hydrophobic antibiotic, to other microbes in membrane vesicles. Environmental Microbiology, v. 22, n. 2, p. 705–713, 2 fev. 2020.

Chougule, Nanasaheb P.; Bonning, Bryony C. Toxins for Transgenic Resistance to Hemipteran Pests. Toxins, v. 4, n. 6, p. 405–429, 4 jun. 2012.

Cuenca, Ana C. P. et al. The Frequency and Spread of a GABA-Gated Chloride Channel Target-Site Mutation and Its Impact on the Efficacy of Ethiprole Against Neotropical Brown Stink Bug, Euschistus heros (Hemiptera: Pentatomidae). Insects, v. 16, n. 4, p. 422, 17 abr. 2025.

Devescovi, Giulia et al. Negative Regulation of Violacein Biosynthesis in Chromobacterium violaceum. Frontiers in Microbiology, v. 8, 7 mar. 2017.

Feng, Zhenhuan et al. DEP2: an upgraded comprehensive analysis toolkit for quantitative proteomics data. Bioinformatics, v. 39, n. 8, 1 ago. 2023.

Ferreira Agüero, Marcos A.; Cremonez, Paulo S. G.; Neves, Pedro M. O. J. Insect Growth Disruptors Cause Mouthpart Malformations, Inhibition of Feeding, and Mortality in the Neotropical Brown Stink Bug Euschistus heros (Hemiptera: Pentatomidae). Journal of Agricultural Science, v. 15, n. 2, p. 40, 15 jan. 2023.

Galdino, Anna Clara M., et al. Disarming Pseudomonas aeruginosa Virulence by the Inhibitory Action of 1,10-Phenanthroline-5,6-Dione-Based Compounds: Elastase B (LasB) as a Chemotherapeutic Target. Frontiers in Microbiology, v. 10, 2 ago. 2019.

González, Esperanza; Falcón-Pérez, Juan Manuel. Expanding Horizons: Next-Generation and Interdisciplinary Advances in the Applications of Extracellular Vesicles. Journal of Extracellular Biology, v. 4, n. 12, 14 dez. 2025.

Jorge, Genesy Pérez et al. Violacein-Loaded Outer Membrane Vesicles from *Salmonella enterica* Exhibit Potent Anti-Melanoma Activity *in Vitro* and *in Vivo*. ACS Biomaterials Science & Engineering, v. 11, n. 10, p. 6166–6184, 13 out. 2025.

Khandelwal, Puneet; Banerjee-Bhatnagar, Nirupama. Insecticidal Activity Associated with the Outer Membrane Vesicles of *Xenorhabdus nematophilus*. Applied and Environmental Microbiology, v. 69, n. 4, p. 2032–2037, abr. 2003.

Kim, Ji Hyun et al. Gram-negative and Gram-positive bacterial extracellular vesicles. Seminars in Cell & Developmental Biology, v. 40, p. 97–104, abr. 2015.

Kowalska, Patrycja et al. Extracellular vesicles of Janthinobacterium lividum as violacein carriers in melanoma cell treatment. Applied Microbiology and Biotechnology, v. 108, n. 1, p. 529, 5 dez. 2024.

Kumar, Pradeep et al. Bacillus thuringiensis as microbial biopesticide: uses and application for sustainable agriculture. Egyptian Journal of Biological Pest Control, v. 31, n. 1, p. 95, 19 dez. 2021.

Li, Huarong; Chougule, Nanasaheb P.; Bonning, Bryony C. Interaction of the Bacillus thuringiensis delta endotoxins Cry1Ac and Cry3Aa with the gut of the pea aphid, Acyrthosiphon pisum (Harris). Journal of Invertebrate Pathology, v. 107, n. 1, p. 69–78, maio 2011.

Martin, Phyllis A. W. et al. Chromobacterium subtsugae sp. nov., a betaproteobacterium toxic to Colorado potato beetle and other insect pests. International Journal of Systematic and Evolutionary Microbiology, v. 57, n. 5, p. 993–999, 1 maio 2007.

Orench-Rivera, Nichole; Kuehn, Meta J. Differential Packaging Into Outer Membrane Vesicles Upon Oxidative Stress Reveals a General Mechanism for Cargo Selectivity. Frontiers in Microbiology, v. 12, 2 jul. 2021.

Patel, Dhruvin H. et al. The Crystal Structure of the Domain of Unknown Function 1480 (DUF1480) From *Klebsiella pneumoniae*. Proteins: Structure, Function, and Bioinformatics, v. 93, n. 3, p. 569–574, 26 mar. 2025.

Pozebon, Henrique et al. Arthropod Invasions Versus Soybean Production in Brazil: A Review. Journal of Economic Entomology, v. 113, n. 4, p. 1591–1608, 13 ago. 2020.

Rausch, Michael A. et al. Modification of Cry4Aa toward Improved Toxin Processing in the Gut of the Pea Aphid, Acyrthosiphon pisum. PLOS One, v. 11, n. 5, p. e0155466, 12 maio 2016.

Remans, Simon et al. Extracellular Vesicles in Arthropods: Biogenesis, Functions, Isolation Methods and Applications. Journal of Extracellular Vesicles, v. 14, n. 9, 3 set. 2025.

Saldanha, Alan Valdir et al. The first extensive analysis of species composition and abundance of stink bugs (Hemiptera: Pentatomidae) on soybean crops in Brazil. Pest Management Science, v. 80, n. 8, p. 3945–3956, 9 ago. 2024.

Sartorio, Mariana G. et al. Bacterial Outer Membrane Vesicles: From Discovery to Applications. Annual Review of Microbiology, v. 75, n. 1, p. 609–630, 8 out. 2021.

Scaramussa, Simone Aparecida de Lima; Ishimoto, Caroline Kie; Bicas, Juliano Lemos. Production, downstream, stability, bioactivities, and applications studies of violacein and related compounds. World Journal of Microbiology and Biotechnology, v. 41, n. 7, p. 226, 25 jul. 2025.

Silva, Paula G. et al. Effect of Insect Growth Regulator Insecticides Novaluron, Teflubenzuron and Lufenuron on the Morphology and Physiology of Euschistus heros. Journal of Agricultural Science, v. 15, n. 7, p. 44, 15 jun. 2023.

Tahir, Muhammad et al. Phosphoproteomic Analysis of Rat Neutrophils Shows the Effect of Intestinal Ischemia/Reperfusion and Preconditioning on Kinases and Phosphatases. International Journal of Molecular Sciences, v. 21, n. 16, p. 5799, 13 ago. 2020.

Tessmer, Magda Andréia et al. Histology of Damage Caused by Euschistus heros (F.) Nymphs in Soybean Pods and Seeds. Neotropical Entomology, v. 51, n. 1, p. 112–121, 21 fev. 2022.

Toyofuku, Masanori et al. Composition and functions of bacterial membrane vesicles. Nature Reviews Microbiology, v. 21, n. 7, p. 415–430, 17 jul. 2023.

Turnbull, Lynne et al. Explosive cell lysis as a mechanism for the biogenesis of bacterial membrane vesicles and biofilms. Nature Communications, v. 7, n. 1, p. 11220, 14 abr. 2016.

Welsh, Joshua A. et al. Minimal information for studies of extracellular vesicles (MISEV2023): From basic to advanced approaches. Journal of Extracellular Vesicles, v. 13, n. 2, 7 fev. 2024.

Wiśniewski, Jacek R. et al. Universal sample preparation method for proteome analysis. Nature Methods, v. 6, n. 5, p. 359–362, 19 maio 2009.

Xie, Junhua et al. The tremendous biomedical potential of bacterial extracellular vesicles. Trends in Biotechnology, v. 40, n. 10, p. 1173–1194, out. 2022.

Zannis-Peyrot, Timothée et al. Extracellular Vesicles of a Phytobeneficial Bacterium Trigger Distinct Systemic Response in Plant. Environmental Microbiology, v. 27, n. 7, 7 jul. 2025.

Zannis-Peyrot, Timothée et al. Toward a better understanding of the ecological roles of extracellular vesicles from plant-associated bacteria. Applied and Environmental Microbiology, 4 nov. 2025.

Zawilak-Pawlik, Anna et al. Establishment of serine protease htrA mutants in Helicobacter pylori is associated with secA mutations. Scientific Reports, v. 9, n. 1, p. 11794, 13 ago. 2019.

Zhang, Yizhuo et al. Vegetative Insecticidal Protein Vip3Aa Is Transported via Membrane Vesicles in Bacillus thuringiensis BMB171. Toxins, v. 14, n. 7, p. 480, 13 jul. 2022.

Zhou, Jun et al. Bacterial Outer Membrane Vesicles: From Physics to Clinical. MedComm – Biomaterials and Applications, v. 4, n. 2, 7 jun. 2025.

Zhou, Xinghong et al. Ginger Extract Decreases Susceptibility to Dextran Sulfate Sodium-Induced Colitis in Mice Following Early Antibiotic Exposure. Frontiers in Medicine, v. 8, 5 jan. 2022.

