## Supplementary data for "Bacterial Extracellular Vesicles from *Chromobacterium subtsugae* and *Bacillus thuringiensis* as Cell-Free Bioinsecticidal Nanocarriers Against the Soybean Pest *Euschistus heros*"

### AFFILIATIONS

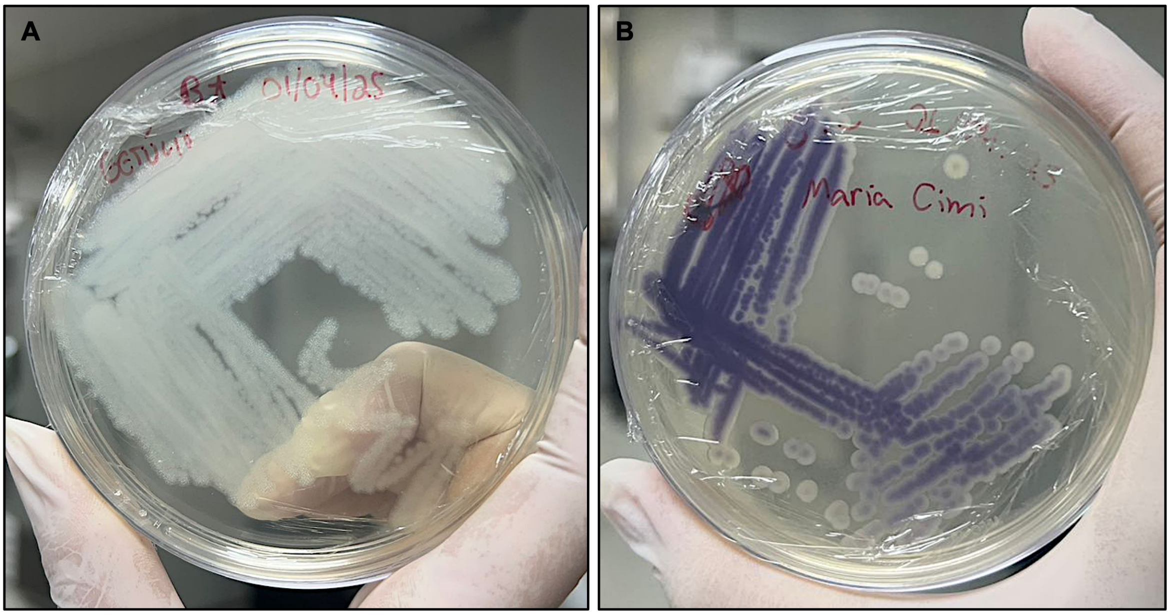


*Bacillus thuringiensis (Bt)* *Chromobacterium subtsugae (Cs)*

**Supplementary Figure 1. Colony morphology of bacterial strains cultured on Tryptic Soy Agar.** (A) *Bacillus thuringiensis* colonies on TSA. (B) *Chromobacterium subtsugae* colonies on TSA. Both strains were cultured under standard conditions (28 °C for *C. subtsugae*, 30 °C for *B. thuringiensis*) for 72 h prior to photography.

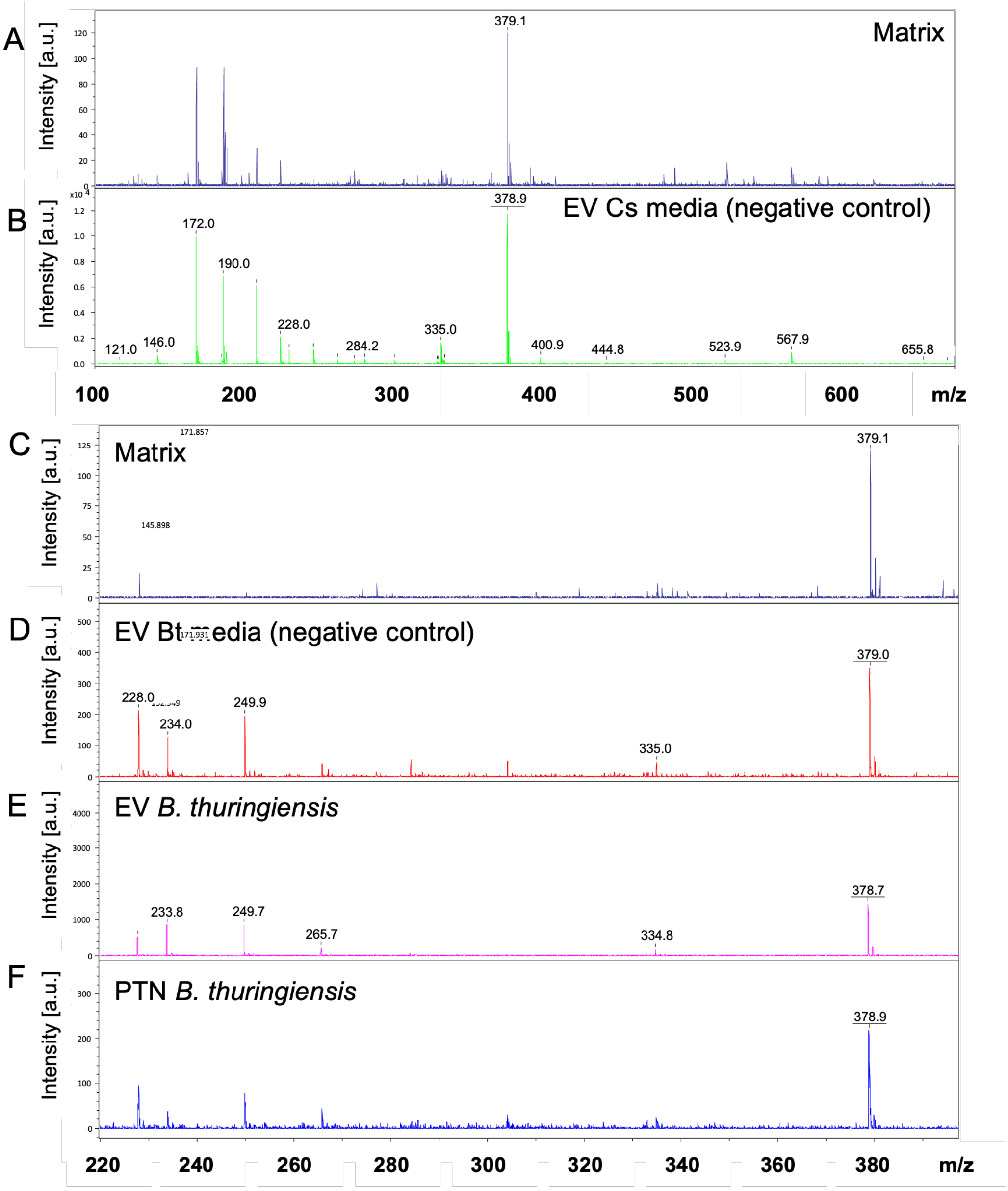


**Supplementary Figure 2. MALDI-TOF mass spectrometry analysis of bacterial extracellular vesicles and soluble protein fractions.** (A) MALDI matrix control spectrum (α-cyano-4-hydroxycinnamic acid) acquired in positive reflector mode, m/z range 100–2000. (B) Mass spectrum of *C. subtsugae* EV culture-media negative control (uninoculated medium), m/z range 100–2000. (C) MALDI matrix control spectrum, m/z range 220–380. (D) Mass spectrum of *Bacillus thuringiensis* EV culture-media negative control (uninoculated medium), m/z range 220–380. (E) Mass spectrum of purified *B. thuringiensis* extracellular vesicles, m/z range 220–380. (F) Mass spectrum of the *B. thuringiensis* soluble protein fraction (PTN), m/z range 220–380. All samples were analyzed on an Autoflex Speed mass spectrometer with external calibration using Peptide Calibration Standard II, in positive reflector mode with 500 laser shots per spot, in triplicate technical replicates. Data processing included baseline subtraction, Savitzky–Golay smoothing (window size 11), and SNAP peak detection (signal-to-noise > 3).

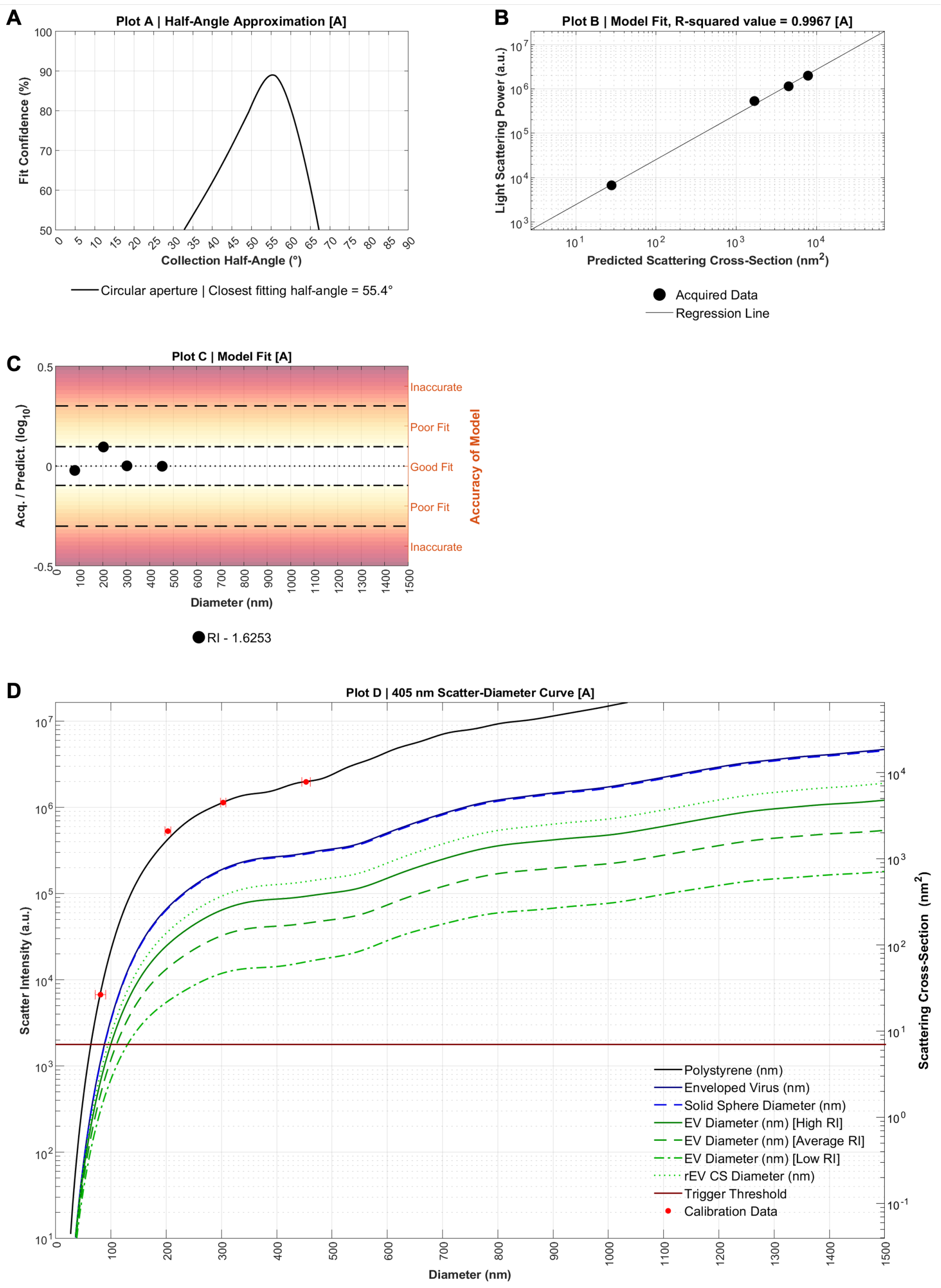


**Supplementary Figure 3. Calibration and validation of nano-flow cytometry using NanoVis beads.** (A) Half-angle approximation plot showing fit confidence as a function of collection half-angle, with circular aperture geometry and a closest-fitting half-angle of 55.4°. (B) Model-fit plot showing the correlation between light-scattering power and predicted scattering cross-section ($R^2$ = 0.9967). (C) Model-fit accuracy across the 100–1500 nm diameter range; the refractive index of the polystyrene calibration standard was RI = 1.6253, with individual points indicating measured calibration particles. (D) 405 nm scatter–diameter curve showing theoretical scattering cross-section versus particle diameter for reference standards, including polystyrene nanoparticles (black solid line), enveloped virus particles (dark-blue solid line), and a solid-sphere model (blue dashed line), with overlaid theoretical curves for extracellular vesicles at high, average, and low refractive index (green shades) and the trigger threshold indicated by red markers. All measurements were acquired on a CytoFLEX S flow cytometer with light scatter calibrated using NanoVis beads (Beckman Coulter).

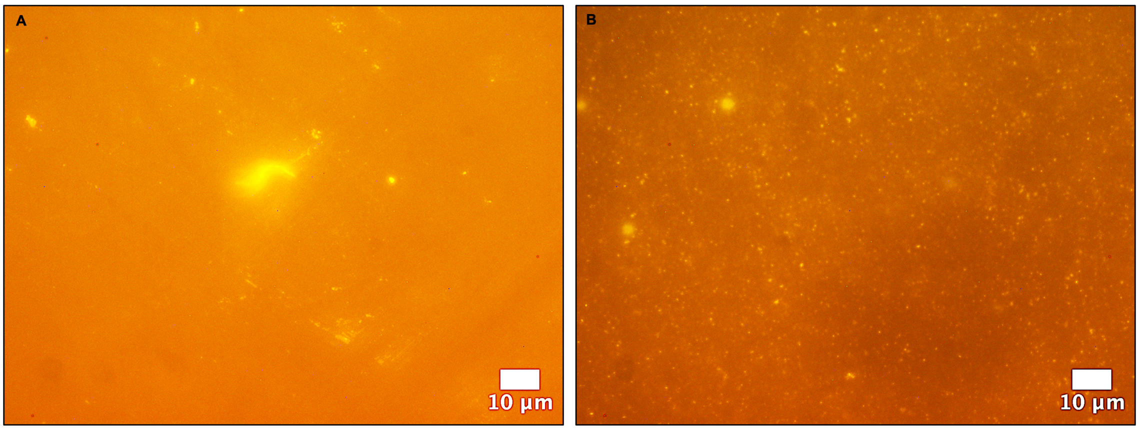


EV from bug media (negative control) bEVs from *C. subtsugae* in bug media

*C. subtsugae*

**Supplementary Figure 4. Fluorescence microscopy of CellMask Deep Red-stained *Chromobacterium subtsugae* extracellular vesicles.** (A) Negative control: uninoculated media processed through the complete bEV isolation workflow (differential centrifugation, 0.22 µm filtration, and SEC) and stained with CellMask Deep Red, showing near-absence of fluorescent puncta. (B) Purified *C. subtsugae* bEVs stained with CellMask Deep Red, showing a dense, uniform field of fluorescent puncta consistent with membrane-labeled nanoparticles. Samples of 5 µL were placed on glass slides with coverslips and imaged on a Zeiss Axio Imager.M2 using a 100× objective and Texas Red filter set (excitation ~560 nm, emission ~630 nm); although the Texas Red filter provides suboptimal excitation for CellMask Deep Red (designed for far-red excitation at ~649 nm), sufficient signal was detected to confirm membrane labeling of the purified bEV fraction. Images are displayed in pseudocolor. Because individual bEVs (~130 nm mean diameter) fall below the optical diffraction limit, each fluorescent punctum represents a diffraction-limited signal from one or more vesicles rather than a resolved individual particle. Scale bar = 10 µm.

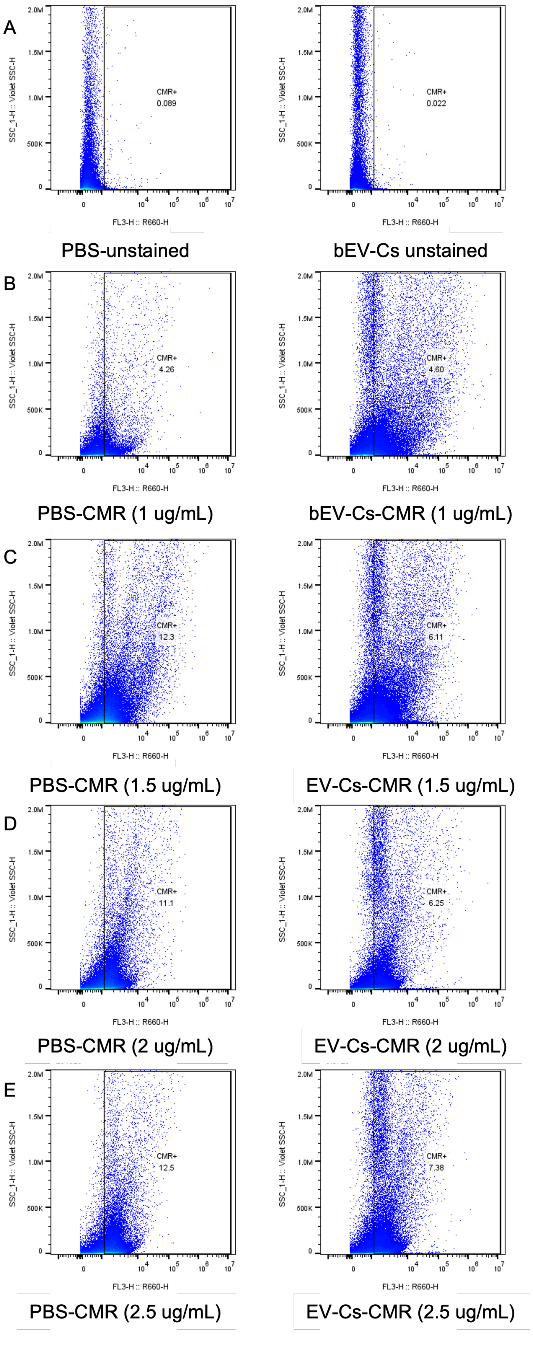


**Supplementary Figure 5. Nano-flow cytometry of unwashed *Chromobacterium subtsugae* extracellular vesicles stained with increasing concentrations of CellMask Deep Red.** Dot plots show side scatter (SSC-H) versus red fluorescence (FL3-H, R660-H detector) for unwashed *C. subtsugae* bEVs stained with CellMask Deep Red (CMR) at increasing concentrations. (A) Unstained controls: PBS only (left) and unstained bEV-Cs (right). (B) PBS and bEV-Cs stained with 1 µg/mL CMR. (C) PBS and bEV-Cs stained with 1.5 µg/mL CMR. (D) PBS and bEV-Cs stained with 2 µg/mL CMR. (E) PBS and bEV-Cs stained with 2.5 µg/mL CMR. Measurements were acquired on a CytoFLEX S flow cytometer with 638 nm excitation. $CMR^+$ gates indicate fluorescently labeled particles. Acquisition: 10,000–50,000 events per sample; doublet discrimination enabled; threshold set to exclude debris.

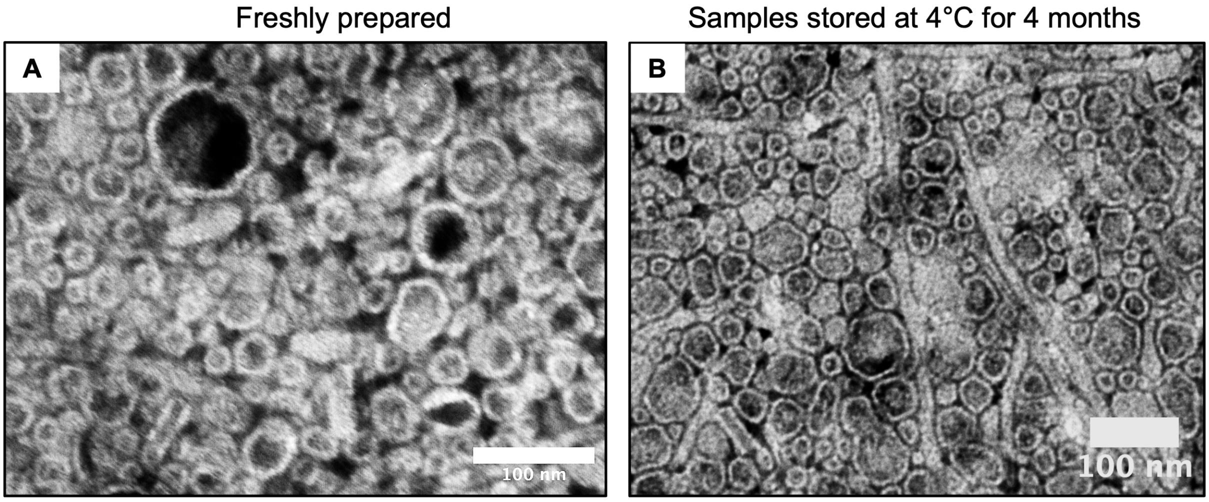


**Supplementary Figure 6. Transmission electron microscopy analysis of *Chromobacterium subtsugae* extracellular vesicle stability.** Negative-stain TEM micrographs of *C. subtsugae* bEVs acquired on a JEOL JEM-1010 microscope at 80 kV. (A) Freshly prepared *C. subtsugae* bEV sample. (B) *C. subtsugae* bEV sample stored at 4 °C for 4 months. Samples were prepared by adsorbing 5 µL of bEV suspension onto Formvar/carbon-coated copper grids for 2 min, washing three times with deionized water, and negatively staining with 1% uranyl acetate for 1 min. Scale bar = 100 nm (both panels).

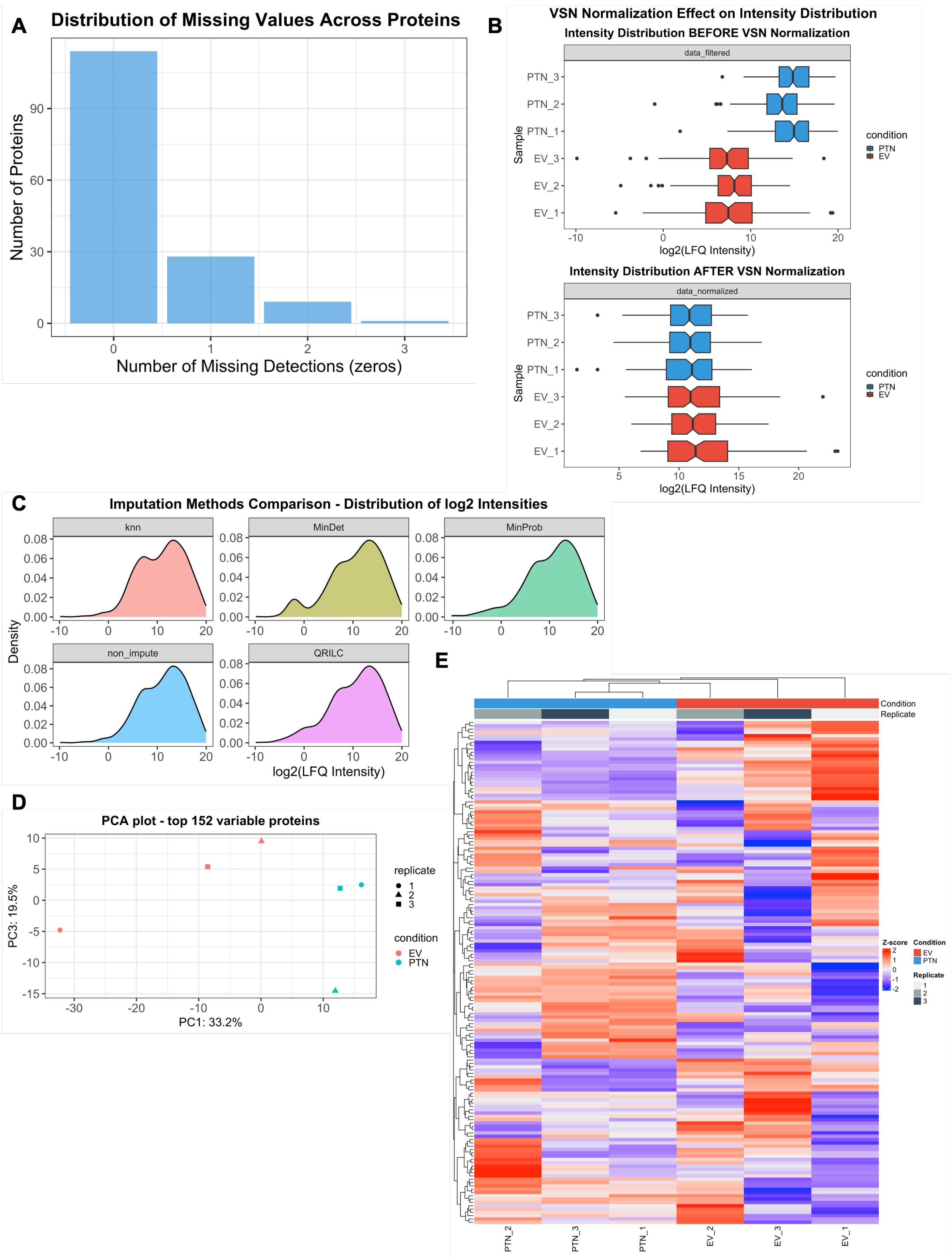


**Supplementary Figure 7. Quality control and statistical analysis of *Bacillus thuringiensis* extracellular vesicle proteomics.** (A) Distribution of missing values across all detected proteins (number of proteins with 0, 1, 2, or 3 missing detections across replicates). (B) Effect of variance-stabilizing normalization (VSN) on intensity distribution: upper panel, before normalization (data_filtered) for soluble protein (PTN, blue) and extracellular vesicle

(EV, red) samples; lower panel, after VSN normalization (data_normalized). (C) Comparison of five imputation methods for missing values — k-nearest neighbors (knn), MinDet, MinProb, non-impute, and QRILC — showing the resulting distributions of $\log_2$ label-free quantitation (LFQ) intensities; the MinProb method was selected for downstream analysis. (D) Principal component analysis (PCA) of the top 152 variable proteins, coloured by condition (EV, red; PTN, teal) and marked by replicate (1, 2, 3); PC1 and PC3 account for 33.2% and 19.5% of variance, respectively. (E) Hierarchical clustering heatmap of normalized $\log_2$ intensities for all detected proteins across samples, organized by condition (EV and PTN) and replicate, with colour indicating relative abundance (blue, low; red, high). Protein identification and quantification used PEAKS Studio v7.0 and Progenesis QI v1.0; normalization, imputation, and the multivariate analyses shown here were performed in R using the DEP2 package.

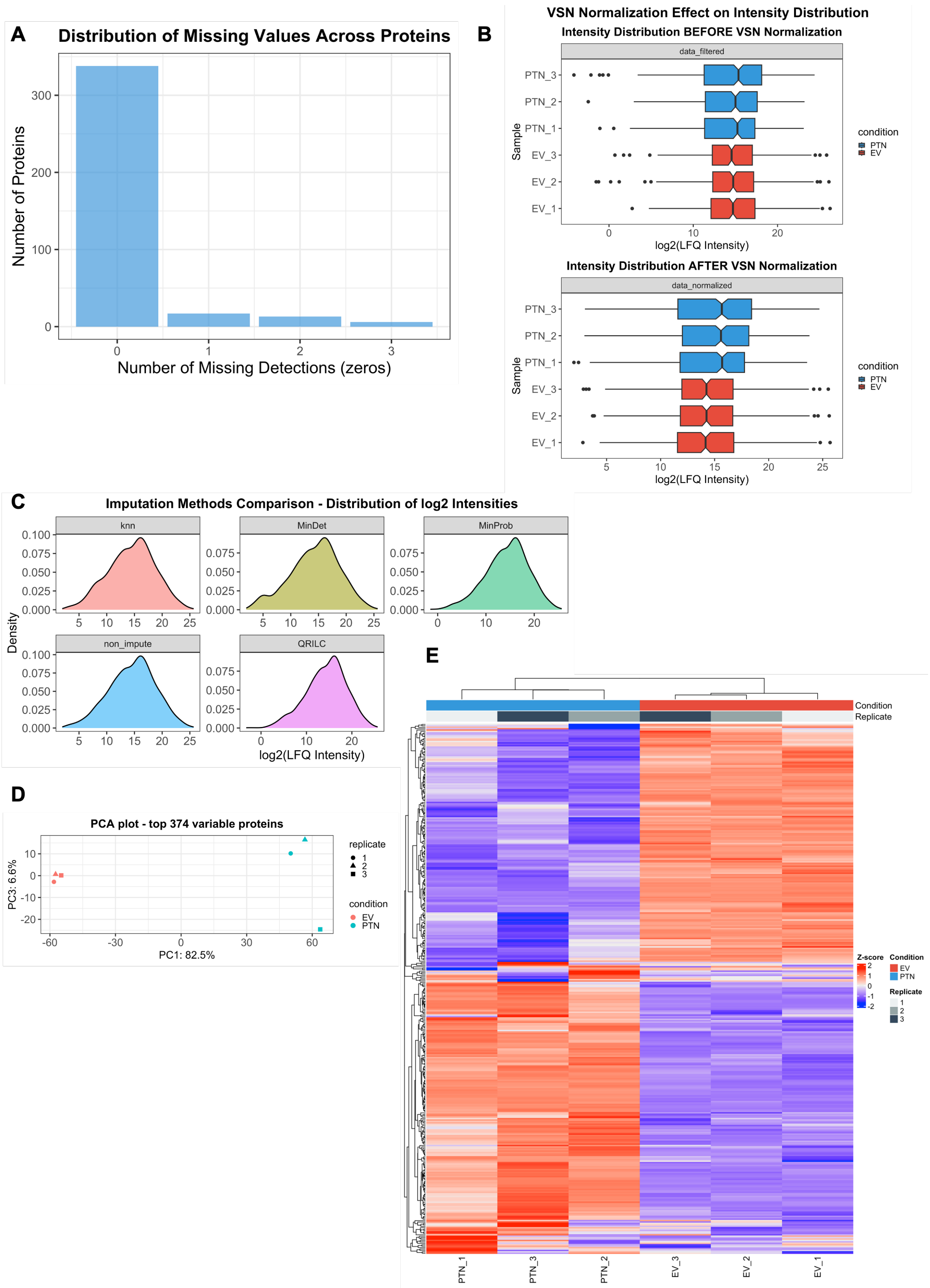


**Supplementary Figure 8. Quality control and statistical analysis of *Chromobacterium subtsugae* extracellular vesicle proteomics.** (A) Distribution of missing values across all detected proteins (number of proteins with 0, 1, 2, or 3 missing detections across replicates). (B) Effect of variance-stabilizing normalization (VSN) on intensity distribution: upper panel, before normalization (data_filtered) for soluble protein (PTN, blue) and extracellular vesicle

(EV, red) samples; lower panel, after VSN normalization (data_normalized). (C) Comparison of five imputation methods for missing values — k-nearest neighbors (knn), MinDet, MinProb, non-impute, and QRILC — showing the resulting distributions of $\log_2$ label-free quantitation (LFQ) intensities; the MinProb method was selected for downstream analysis. (D) Principal component analysis (PCA) of the top 374 variable proteins, coloured by condition (EV, red; PTN, teal) and marked by replicate (1, 2, 3); PC1 and PC3 account for 82.5% and 6.6% of variance, respectively. (E) Hierarchical clustering heatmap of normalized $\log_2$ intensities for all detected proteins across samples, organized by condition (EV and PTN) and replicate, with colour indicating relative abundance (blue, low; red, high). Protein identification and quantification used PEAKS Studio v7.0 and Progenesis QI v1.0; normalization, imputation, and the multivariate analyses shown here were performed in R using the DEP2 package.
